# Expectation-Maximization enables Phylogenetic Dating under a Categorical Rate Model

**DOI:** 10.1101/2022.10.06.511147

**Authors:** Uyen Mai, Eduardo Charvel, Siavash Mirarab

## Abstract

Dating phylogenetic trees to obtain branch lengths in time unit is essential for many downstream applications but has remained challenging. Dating requires inferring substitution rates that can change across the tree. While we can assume to have information about a small subset of nodes from the fossil record or sampling times (for fast-evolving organisms), inferring the ages of the other nodes essentially requires extrapolation and interpolation. Assuming a clock model that defines a distribution over rates, we can formulate dating as a constrained maximum likelihood (ML) estimation problem. While ML dating methods exist, their accuracy degrades in the face of model misspecification where the assumed parametric statistical clock model vastly differs from the true distribution. Notably, existing methods tend to assume rigid, often unimodal rate distributions. A second challenge is that the likelihood function involves an integral over the continuous domain of the rates and often leads to difficult non-convex optimization problems. To tackle these two challenges, we propose a new method called Molecular Dating using Categorical-models (MD-Cat). MD-Cat uses a categorical model of rates inspired by non-parametric statistics and can approximate a large family of models by discretizing the rate distribution into k categories. Under this model, we can use the Expectation-Maximization (EM) algorithm to co-estimate rate categories and branch lengths time units. Our model has fewer assumptions about the true clock model than parametric models such as Gamma or LogNormal distribution. Our results on two simulated and real datasets of Angiosperms and HIV and a wide selection of rate distributions show that MD-Cat is often more accurate than the alternatives, especially on datasets with nonmodal or multimodal clock models.

**Code availability:** The MD-Cat software is available at https://github.com/uym2/MD-Cat.

**Data availability:** Data are available on Github https://github.com/uym2/MD-Cat-paper and Dryad https://doi.org/10.5061/dryad.pk0p2ngs0.

## 1 Introduction

Rates of evolution can change through time, creating a major challenge for dating phylogenetic trees. Since time and substitution rates are inseparable from sequence data, dating relies on pre-specified ancestral divergence times (e.g., calibration points obtained from radiometric dating fossils) or sampling times of the tips (available for phylodynamic data) to estimate the rates of evolution. However, even with such external information, extrapolating from a few known divergence times to all nodes essentially requires assumptions about how the rates may or may not have changed. Ideally, we need to know a *molecular clock* model, giving us a way to assign probabilities to substitution rates throughout the tree.

The simple strict clock model by Zuckerkandl and Pauling (1962) is insufficient, especially at long evolutionary time horizons where substitution rates have had enough time to change (Bromham and Penny, 2003; Kumar, 2005). As a result, various models have been developed to allow the rates to change across the tree, relaxing the strict clock assumption. Uncorrelated models assume that rates are drawn i.i.d. for each branch from a parametric distribution, such as exponential, gamma, or lognormal (Aris-Brosou and Yang, 2002), while autocorrelated models allow the rate to evolve on the tree and capture the correlation between parent and child branches. Computational methods for molecular dating have been developed for both uncorrelated and autocorrelated clock models (see Ho and Duchêne, 2014; Kumar and Hedges, 2016; Rutschmann, 2006; Sanderson, 1998). In particular, Bayesian methods (e.g., Drummond and Rambaut, 2007; Guindon, 2010; Heath, 2012; Rannala and Yang, 2007; Thorne and Kishino, 2002; Thorne et al., 1998) have been popular, perhaps because they can incorporate complex rate models thanks to the use of MCMC sampling. Bayesian methods can infer a dated tree directly from sequence data, jointly estimating the topology, branch lengths, rates, and thus divergence times. The availability of priors also enables users to control the amount of rate heterogeneity across the tree, which can be helpful when sequence data are less informative. Nevertheless, there has been debate around the merit of Bayesian methods (Beaulieu et al., 2015; Wertheim et al., 2010). One practical hurdle is that choosing the model and the prior is challenging. Moreover, the computational burden of the MCMC process limits its scalability.

Maximum likelihood (ML) methods promise to improve scalability compared to MCMC (e.g., Langley and Fitch, 1974; To et al., 2016; Volz and Frost, 2017). These methods assume a statistical model for the rates and seek parameters that maximize the likelihood. ML-based methods usually do not work directly on sequence data but take as input a tree with branch lengths in the unit of substitutions per site (SU), inferred from sequence data; these methods then convert the SU branch lengths to time units using calibration points (Bayesian methods using this two-step approach have also been proposed; e.g., Didelot et al. (2018)). Although they are more efficient than Bayesian methods and do not need a prior distribution, ML-based methods have their own limitations. The simplest ML methods assume a strict clock model and use either a Poisson (Langley and Fitch, 1974) or Gaussian (To et al., 2016) distribution to model the uncertainties of the estimated branch lengths in substitutions per site. These ML methods, despite being fast, are less commonly used because of the strict clock model they assume. More sophisticated ML methods, such as TreeDater (Volz and Frost, 2017) and TreeTime (Sagulenko et al., 2018), use a Gamma or LogNormal distribution to model the molecular clock. Because they assume a parametric – and often unimodal – clock model, these methods may not be robust under complex scenarios with local clocks or sudden rate changes. In addition, the substitution rates of these models are continuous latent variables, so the likelihood function contains an intractable integral, making the likelihood function difficult to optimize. Therefore, some ML-based methods (e.g., Volz and Frost, 2017) depend on heuristic algorithms to iteratively optimize their likelihood functions. These heuristic algorithms lack theoretical guarantees to reach a local (or global) optimum, a monotonic improvement of the likelihood function, or convergence.

Recognizing the difficulties of relying on a specific clock model, many authors, including us (Mai and Mirarab, 2020), have proposed non-parametric methods for dating (e.g., Sanderson, 1997, 2002; Tamura et al., 2012, 2018; Xia and Yang, 2011). These methods formulate dating as an optimization problem — often in a least-square form — to optimize a predefined objective function without an explicit parametric model of the clock. However, the objective functions used in these methods come from implicit assumptions about the rate distribution. For example, most methods implicitly assume that rates are distributed following a unimodal distribution; that is, the rate of each branch is centered either around a global rate (Mai and Mirarab, 2020; Xia and Yang, 2011) or the rate of its parent or sibling (Britton et al., 2007; Sanderson, 1997; Tamura et al., 2012). Incidentally, minimizing the residual sum-of-square often winds up being the ML solution under specific (often unimodal) models; e.g., our attempt at developing the non-parametric method wLogDate (Mai and Mirarab, 2020) produced a method that is the ML estimate under a set of (unimodal) LogNormal distributions. Thus, although non-parametric methods are fast and show signs of robustness under multiple clock models, we postulate (and test) that these non-parametric methods are inaccurate when the true clock model is multimodal or is a combination of multiple local clocks.

To increase the robustness of statistical models, many authors have turned to categorical distributions (often called CAT) for modeling various evolutionary processes. CAT models have been adopted to approximate the Gamma model for rate heterogeneity across sequence sites (Felsenstein, 2004; Nguyen et al., 2015; Stamatakis, 2006). They have also been used extensively in modeling the heterogeneity of substitution matrices and base frequencies across sites (e.g., Foster, 2004; Huelsenbeck et al., 2006; Lartillot and Philippe, 2004) and heterotachy (e.g., Blanquart and Lartillot, 2008). More relevant to this work are the discrete-rate models of substitution rate heterogeneity across the tree (Fourment and Holmes, 2014) and Bayesian rate priors based on categorical rates (Heath et al., 2012). These attempts have shown a path towards robust models that allow flexibility in the underlying distribution of model parameters. The present work is motivated by these attempts but addresses the persistent computational challenges (we return to a comparison of our approach to these in the Discussion section).

In this paper, we introduce a new ML method called Molecular Dating with Categorical rates (MD-Cat). Unlike most ML-based methods, we use a categorical distribution to approximate the unknown continuous clock model (Fig. 1). The model is defined as a set of *k* rate categories, each captured by a free parameter, and branch rates are drawn from the resulting discrete distribution. Although the rate categories are discrete, the model has the power to approximate a continuous clock model if *k* is large and there is enough data to fit the model – a notion that Höhna et al. (2019) had earlier used to propose a cladogenesis model with varying diversification rates. We use the Expectation-Maximization (EM) algorithm to maximize the likelihood function associated with this model, where the *k* rate categories and branch lengths in time units are modeled as unknown parameters and co-estimated. We show that both the E-step and M-step of our EM algorithm can be computed efficiently, and the algorithm is guaranteed to converge. We compare MD-Cat to state-of-the-art methods on a diverse set of simulations, including multimodal models and local clocks.

**Figure 1.**
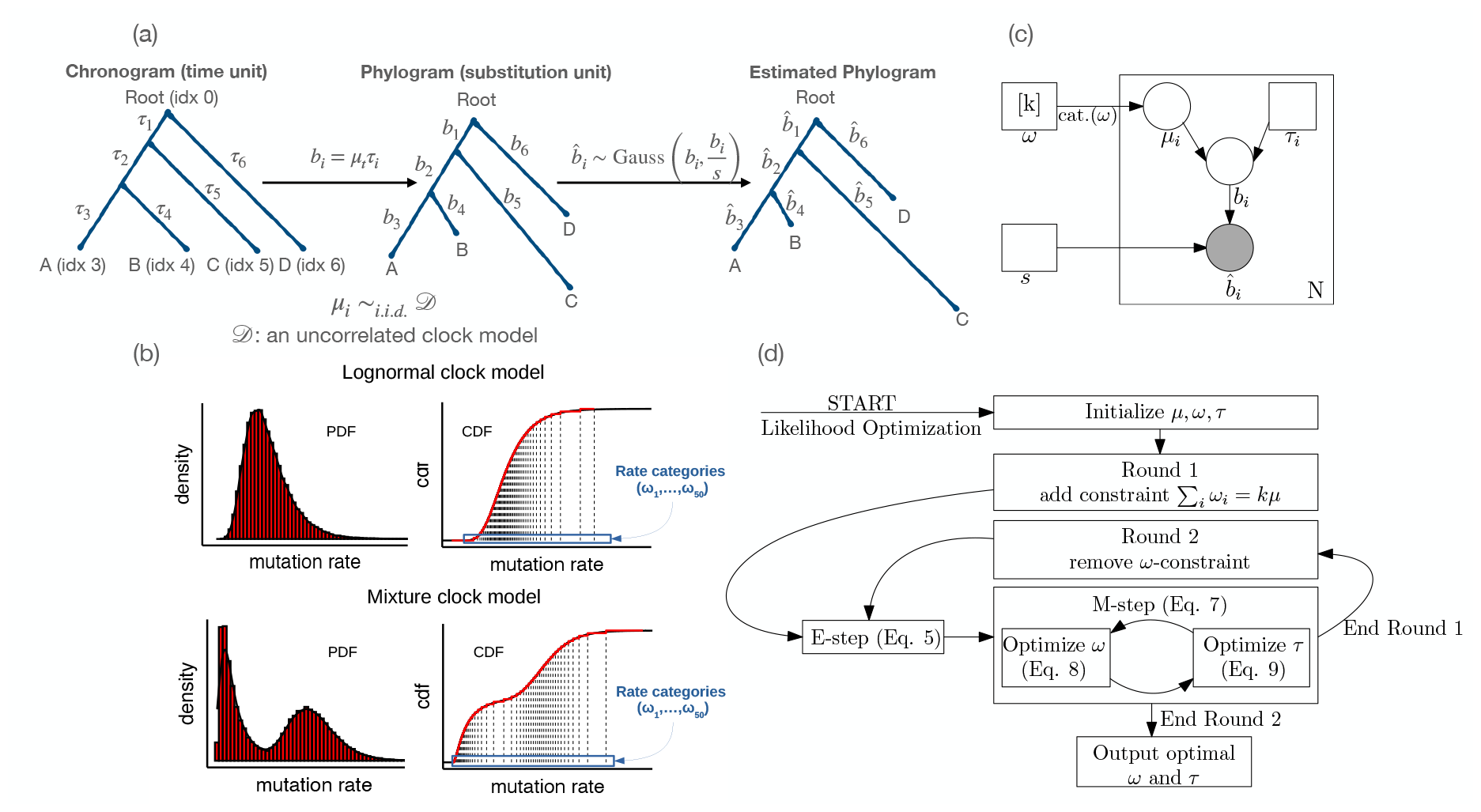
(a) The generative model. (b) Examples of clock models and their categorical approximation. The approximation can be visualized on the PDF as a histogram and on CDF as a step function (shown with *k* = 50). Because we give the same weight to each rate category, the step function has of equal vertical steps. Rate categories are identified by the projection of the steps onto the x-axis of the CDF plot. (c) The hierarchical categorical clock model. We use the plate notation where squares denote parameters, circles denote random variables, and solid circles denote observed data. The number inside a plate (e.g. *N*) denotes the number of repeats of each random variable inside that plate. (d) Steps of the optimization algorithm.

## 2 Methods

### 2.1 Notations and Model

For a given binary tree *T* with *n* leaves and *N* = 2*n −* 2 branches, we give each of the *n −* 1 internal nodes of *T* a unique index in {0, …, *n −* 2} (reserving 0 for the root) and give each of the *n* leaf nodes a unique index in *{n −* 1, …, *N*}. We denote the parent of node *i* as *par*(*i*), the left and right children of node *i* as *c*_*l*_(*i*) and *c*_*r*_(*i*), and the edge connecting *i* and *par*(*i*) as *e*_*i*_ for *i* ∈ {1, …, *N*}. The length of *e*_*i*_ is specified in either SU or time unit. Let *t*_*i*_ denote the divergence time of node *i* (i.e. the time when species *i* diverges into *c*_*l*_(*i*) and *c*_*r*_(*i*)). This time is measured from some arbitrary but fixed origin (e.g., present day) and can be either positive (after origin) or negative (before origin). Then *τ*_*i*_ = *t*_*i*_ *− t*_*par*(*i*)_ is the length of *e*_*i*_ in time unit (Fig. 1a). As a shorthand, we combine all the *τ*_*i*_ values into a vector *τ* = [*τ*_1_, *τ*_2_ …, *τ*_*N*_ ]. Similarly, let *b*_*i*_ be the length of *e*_*i*_ in SU, which is the *expected number of substitutions per sequence per site* occurred on edge *e*_*i*_ and let **b** = [*b*_1_, *b*_2_, …, *b*_*N*_ ]. We assume that the substitution rate can change only at species divergence times and let *µ*_*i*_ = *b*_*i*_*/τ*_*i*_ denote the rate along branch *e*_*i*_. Finally, we let *s* denote the alignment length.

#### Model

Sequences of length *s* evolve down a binary tree *T* with substitution rate *µ*_*i*_ along each edge *e*_*i*_. We assume *µ*_*i*_ are i.i.d following an unknown distribution (e.g., Fig. 1b) and sites of the molecular sequences evolve independently following a homogeneous process such as the GTR model (Tavaré, 1986). We are given an estimate of the topology of *T* and all its SU branch lengths. This estimate can be obtained using standard computational methods (e.g., ML or distance-based methods) and can contain errors. We assume the tree topology is estimated correctly but branch lengths have error. Let *B*_*i*_ be a random variable denoting the estimated SU length of branch *e*_*i*_ and 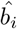 be the estimate of *b*_*i*_; then, 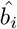 is an observation of *B*_*i*_ (Fig. 1a). The distribution of *B*_*i*_ depends on several factors such as the sequence evolution model, sequence length, and the inference technique. For simplicity, we model *B*_*i*_ similarly to To et al. (2016): let *ϵ*_*i*_ = *B*_*i*_ *− b*_*i*_ be the estimation error and assume 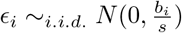, where the variance *b*_*i*_*/s* comes from an approximation of the Poisson model of the number of substitutions per sequence site. Recall that *b*_*i*_ = *µ*_*i*_*τ*_*i*_ and *B*_*i*_ = *b*_*i*_ + *ϵ*_*i*_; thus, 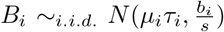. For computational and algorithmic convenience, we approximate the variance 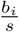 by 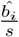 and let 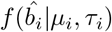 denote the associated Gaussian pdf.

As mentioned previously, the clock model that describes the distribution of *µ* is uncertain. Common choices include Gaussian (Sagulenko et al., 2018; To et al., 2016), Gamma (Volz and Frost, 2017), LogNormal, and Exponential distributions (Drummond et al., 2006). However, there is no guarantee that these distributions, which are unimodal or mode-less, can capture how *µ* changes across a tree. For example, a bimodal or tri-modal set of rates would not be adequately modeled with any of these distributions. Instead of using a parametric continuous distribution, we use an approximation approach using a discrete distribution. We discretize *µ* into *k* categories *ω* = [*ω*_1_, *ω*_2_, …, *ω*_*k*_] each with the same probability mass ^1^*/k*. The *ω*_*i*_s are free parameters, which we will estimate from the data. By adjusting the position of *ω* parameters, this model can approximate any continuous distribution (see Fig. 1b). We use this categorical model to approximate the unknown distribution of *µ*. Putting together all these elements, we obtain a hierarchical model (Fig. 1c) where *τ* and *ω* are parameters, 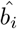 ’s are observations (i.e. data), and both *b*_*i*_’s and *µ*_*i*_’s are latent variables.

#### The linear constraints defined by calibration points

Let *t*_0_ be the *unknown* divergence time at the root of the tree. We incorporate the *p* calibration points *t*_1_, …, *t*_*p*_ as a set of *p* constraints *C*_1_, …, *C*_*p*_ :

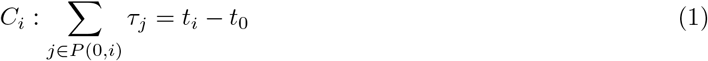

where *P* (*x, y*) denotes the path between two nodes *x* and *y* (referring to by their indices). Recall that the root has index 0, so *P* (0, *i*) is the path from the root to node *i*. To remove *t*_0_ from this set of constraints, we arbitrarily select a constraint *C*_*k*_ and subtract it from the other constraints *C*_*i*_ (*i*≠ *k*) side by side to obtain the set Ψ^(*k*)^ of *p −* 1 linear constraints on *τ* :

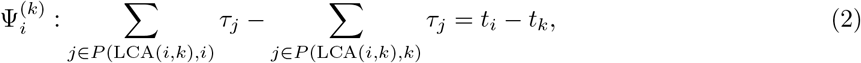

It is easy to see that all the linear constraint sets Ψ^(*k*)^ (*k ∈* [*p*]) are equivalent. Therefore, we use Ψ as a shorthand to refer to any (arbitrarily chosen) among them. We can construct a set Ψ using a bottom-up traversal of the tree, as we have done elsewhere (Mai and Mirarab, 2020).

### 2.2 Maximum likelihood estimation using EM algorithm

Under the categorical model of the rates and the Gaussian model of branch length estimation error, the log-likelihood of 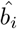 given the parameters *τ* and *ω* is

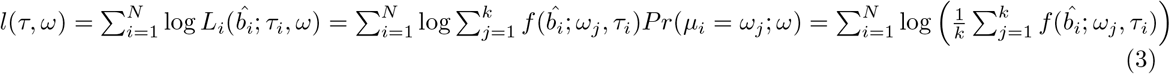

where 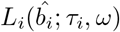 denotes the density of 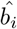 on branch *i* and *f* is the density function of the Gaussian model. Our goal is to find *τ* and *ω* that maximize *l*(*τ, ω*) and satisfy Ψ. Because *l* has a summation inside the log function, it is difficult to optimize it directly. However, thanks to the categorical model of the rates, the latent variables *µ*_*i*_ are discrete, and we can readily apply the EM algorithm (Dempster et al., 1977). We start with an initial *τ* and *ω* (construction of these is described later) and iteratively improve the log-likelihood function by alternating between the E-step and M-step. In the **E-step**, we compute the posterior of the latent variables, setting

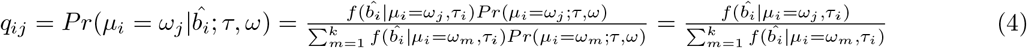

where the second equality holds because 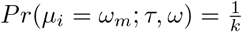 for all *i, m*. Note these values are trivial to compute as *f* is simply the Gaussian pdf. In the **M-step**, we find *ω* and *τ* to maximize

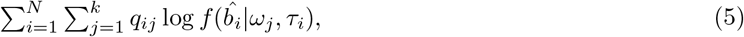

such that the constraints Ψ given in (2) are satisfied. Using the Gaussian pdf and removing constants, we obtain the following equivalent optimization problem:

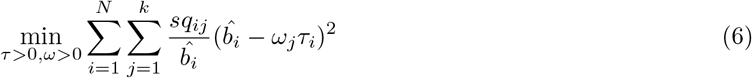

such that Ψ is satisfied. This problem is non-convex and is difficult to solve optimally. However, it is easy to see that the likelihood function is bounded above, so it has a maximum. Therefore, the EM algorithm still converges as long as after every iteration (h) the M-step finds a new point (*τ* ^(*h*+1)^, *ω*^(*h*+1)^) that gives a higher value for Eq. (5) (or equivalently, lower value for Eq. (8)) than that of (*τ* ^(*h*)^, *ω*^(*h*)^). To do that, we start with *ω*^(*h*)^ and *τ* ^(*h*)^ and successively minimize Eq. (8) along the coordinate block of either *ω* or *τ* while fixing the other and iterate until convergence (i.e. block coordinate descent). In other words, let *τ* ^(*h*,1)^ = *τ* ^(*h*)^ and *ω*^(*h*,1)^ = *ω*^(*h*)^; in each iteration (p) of coordinate descent, we find *τ* ^(*h,p*+1)^ and *ω*^(*h,p*+1)^ such that:

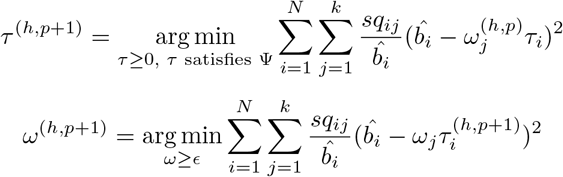

These two problems are instances of the *weighted least-square* optimization that can be solved efficiently using convex programming. Inspired by To et al. (2016), we use the active-set method to solve these two problems. With an efficient use of the Lagrange’s method, the complexity of each iteration of the active-set method is *O*(*N* + *k*) (see section A.1 and Algorithms 1 and 2 in Supplementary). Let *p*^*∗*^ be the iteration when block coordinate descent converged, then we set 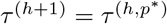 and 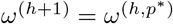

#### Initialization

The EM algorithm can only find local optima, so it needs multiple initial points. To facilitate the search, we first estimate the expected substitution rate *µ*, then discretize the uniform distribution [0, 2*µ*] into *k* equal segments and set 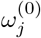 to the middle of the *j*^*th*^ segment. Thus,

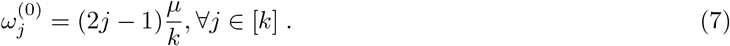

To initialize 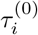, we draw *j* uniformly in [*k*] and set 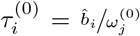. Although this initialization does not guarantee that 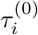 satisfy Ψ, the constraints will be satisfied after the first M-step. We run wLogDate (Mai and Mirarab, 2020) and root-to-tip (RTT) regression (Shankarappa et al., 1999) to get two different estimates of *µ*. We then use each of these two estimates to get *m* different initials for *τ* ^(0)^, for a total of 2*m* initials. We use *m* = 100 by default.

#### Two-round optimization

A difficulty in co-estimating rates and times is that they are inseparable as a product. This same problem occurs in the EM algorithm; in each iteration, *ω* and *τ* can both be scaled up or down by the same factor, only constrained by calibration points. To avoid rapid jumps of *ω* and *τ*, we use a two-step approach (Fig. 1d). We first enforce *ω* to have the mean *µ* (estimated by either wLogDate or RTT, as described above) and run EM until convergence. Then we relax that constraint and let EM continue running to re-estimate the expected *µ*. Thus, for each initial point, we run EM twice. In the first round, the expected substitution rate is fixed to the initial *µ* by adding the constraint 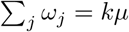 to Ψ. In the second round, we initialize EM with the *ω* and *τ* found in the first round, relax the constraint 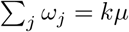, and let the EM algorithm run until convergence. The final solution is the one that gives the lowest value for Eq. (8) among all initial points. We claim the following result:

##### Claim 1.

*The EM algorithm described above monotonically improves the log-likelihood function after each iteration. Furthermore, if the log-likelihood l*(*τ, ω*) *has a maximum value, then the algorithm converges*.

A proof is given in section A.2 in the Supplementary. Consistent with the theory, our empirical tests show that for any initial point, the log-likelihood increases monotonically after each iteration and eventually converges (Fig. S1 in the Supplementary).

#### Support values

To obtain confidence intervals (CI) for our point estimates, we perform bootstrap on *i* the estimated rate categories, as follows. Recall that after the last iteration of the EM algorithm, we obtain the estimated rate categories *ω*^*∗*^. We then perform a user-defined number of parametric bootstrap steps (default: 1000). For each bootstrap replicate *r*, for each branch *i*, we draw a rate 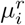 from *ω*^*∗*^. After every branch has been assigned a rate 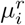, we find *τ* ^*r*^ as follows:

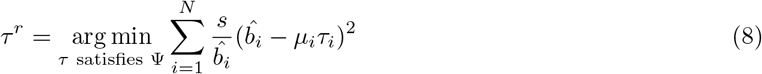

Note that computing *τ* ^*r*^ is very fast because the above problem is a standard weighted least-square optimization. After computing *τ* ^*r*^, we also compute all divergence times 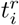 for every node *i*. Having access to all the bootstrap estimates, we compute an empirical *α*-level CI for each estimate *τ*_*i*_ by taking the *α/*2 and 1 *− α/*2 percentiles of 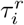 values. We compute the CI for each divergence time *t*_*i*_ similarly.

#### Automatic *k* selection

MD-CAT uses a fixed *k* = 50 by default. At the expense of higher running time, one can use a model selection method such as Akaike information criterion (AIC) or Bayesian information criterion (BIC) to select *k*. Alternatively, if many calibration points are available, we can use a crossvalidation approach to select the *k*, as follows. We create *r* replicate datasets by subsampling a portion *p* of calibration points at random and hiding the rest. For each replicate, we then run MD-CAT using the sampled calibration points with a range of *k* values (e.g., *k* ∈ {2, 5, 10, 25, 50, 100}) and measure the error in estimating the divergence time of the points kept hidden in that replicate. The *k* value with the lowest mean divergence time error is selected for a final run with all the calibration points included. By default, we set *r* = 5 and *p* = 0.8.

### 2.3 Evaluation: datasets

We used a simulated and a real angiosperm dataset and a simulated and a real HIV *env* gene dataset to evaluate MD-Cat in comparison to existing methods, LSD, BEAST, wLogDate, and RelTime.

#### Simulated angiosperms with hybrid rates

Dating the emergence of flowering plants (angiosperms) is a well-studied example of the discrepancy between molecular dating and fossil records. Recent studies using molecular sequences report the estimate of the angiosperm crown age to be around 190-250 Mya (Clarke et al., 2011; Foster et al., 2017; Zeng et al., 2014), i.e., early Jurassic to Triassic, while the oldest fossils date around 140 Mya, leaving a gap of as much as 100 million years between molecular dating and fossil records.

Beaulieu et al. (2015) studied the effect of hybrid substitution rates on the accuracy of dating angiosperms. They simulated a dataset emulating angiosperms where evolutionary rates formed local clocks in groups of clades. Their results showed that Bayesian estimation can perform poorly on such a complex clock model, providing a possible explanation for the gap between molecular dating and fossil records. In their simulations, the true age in all replicates was fixed to 140 mya. They simulated five clock model scenarios with two or three rates across several clades. See supplementary Table S2 and Beaulieu et al. (2015) for details of the five scenarios and note that going from scenario 1 to 5, rates become more varied. The time tree and 100 simulated phylograms for each of these five scenarios were downloaded from the Dryad Repository provided by the authors. We used the provided phylograms to simulate DNA sequences of length 1000 using SeqGen under the GTR model with the shape of the Gamma rate heterogeneity across sites set to *α* = 1. We compared MD-Cat to three other methods: BEAST (Drummond and Rambaut, 2007), wLogDate (Mai and Mirarab, 2020), and RelTime (Tamura et al., 2012). We used the true rooted tree topology and inferred branch lengths in substitutions per site using RAxML (Stamatakis, 2006). We used the 20 species on the clade outside the angiosperms as the outgroup to root the RAxML tree. Because outgroup rooting cannot determine the exact root position on the branch connecting ingroups and outgroups, we removed the outgroups from the tree before dating. As such, five calibration points that belong to the 20 species in the outgroups were discarded, leaving us with 15 calibration points. We dated the tree using MD-Cat, wLogdate, and RelTime, providing each with the exact time of those 15 calibration points. We ran wLogDate with 100 feasible starting points, MD-Cat with *k* = 50 rate categories and 200 initial points, and RelTime with default settings.

We ran BEAST with the topology fixed to the true topology and the same settings as Beaulieu et al. (2015): a LogNormal (LN) prior on rates and the MCMC chain of 30 million. Our ESS for the posterior root age is well above 100 for a majority of the replicates (Supplementary Fig. S2), showing evidence of convergence. In scenario 1, we also tested two other priors, strict molecular clock and random local clock (RLC) (Drummond and Suchard, 2010). We did not run BEAST with these priors on other datasets due to the high running times needed.

#### Simulated HIV phylodynamics

We reused the phylodynamics data of HIV *env* gene simulated by To et al. (2016) but explored many more clock models. To et al. (2016) simulated time trees based on a birth-death model with periodic sampling times. There are four tree models: D995 11 10 (M1) and D995 3 25 (M2) simulate intra-host HIV evolution and D750 11 10 (M3) and D750 3 25 (M4) simulate inter-host evolution. Each tree model has 100 replicates, each with either 110 leaves (M1 and M3) or 75 leaves (M2 and M4). In our earlier work (Mai and Mirarab, 2020), from the given time trees we simulated trees with SU branch length using three clock models: LogNormal, Gamma, and Exponential. In this paper, we augmented this dataset with nine new clock models: uniform distribution, four mixtures of two, three mixtures of three, and a mixture of four LogNormal distributions (Fig. 3a and Supplementary Table S1).

We simulated sequence data using Seqgen under the same settings as To et al. (2016) (sequence length: 1000; DNA evolution model: the F84 model with a gamma distribution for across-sites rate heterogeneity with shape 1.0 and eight rate categories; transition/transversion rate ratio: 2.5; nucleotide frequency of A, C, G, T: 0.35, 0.20, 0.20, 0.25). We compared MD-Cat to LSD (To et al., 2016), wLogDate (Mai and Mirarab, 2020) and BEAST (Drummond and Suchard, 2010). We used the true rooted tree topology. To test LSD, wLogDate, and MD-Cat, we used RAxML (Stamatakis, 2006) to estimate the SU branch lengths from the simulated sequences using the GTRGAMMA model and used each of these methods to infer the time tree. LSD was run with the same settings as the QPD* mode described by To et al. (2016), and wLogDate was run with 100 feasible starting points. MD-Cat was run with *k* = 50 rate categories and 200 initial points. To test BEAST, we used the sequences simulated by SeqGen, fixed the true rooted tree topology, and only inferred node ages. We ran BEAST using the following priors: HKY+G8 model, coalescent with constant population size, and strict-clock (i.e. fixed rate) clock model. We set the length of the MCMC chain to 5 *×* 10^6^ generations, burn-in to 10%, and sampling to every 10^4^ generations.

#### Real angiosperm data

We used MD-Cat to reanalyze the biological angiosperm data by Beaulieu et al. (2015) consisting of DNA sequences of four genes (atpB, rbcL, psbA, and 18S). We reused the rooted tree topology estimated by BEAST from the original study and ran RAxML on the concatenated alignment to estimate the SU branch lengths (fixing the topology). We then used MD-Cat to estimate the divergence times and per-branch substitution rates. We used the same 24 calibration points as Beaulieu et al. (2015), except for the calibration for angiosperm, which is the focal node of the evaluation. Each calibration point was incorporated as a point value (i.e., equality).

#### HIV LANL dataset

We retrieved the alignment of the *env* region for HIV-1 M group, which includes subtypes A–L, from the Los Alamos HIV Sequence Database, LANL (http://www.hiv.lanl.gov/). The dataset includes 4062 sequences sampled from 1981 to 2020, with sampling year provided (328 sequences that have missing data were removed). We randomly subsampled subtypes B and C such that each of them has at most 20 sequences for each distinct year and kept all sequences of the other subtypes. As such, 1104 sequences remained, consisting of 2 subtype A, 208 subtype A1, 4 subtype A2, 447 subtype B, 200 subtype C, 99 subtype D, 41 subtype F1, 13 subtype F2, 79 subtype G, and 11 sequences of subtypes H-L. We used IQTree (with ModelFinder, which selected the GTR+F+I+I+R10 model) to estimate the tree topology and branch lengths, then rooted the tree using root-to-tip regression (Mai et al., 2017). Because two subtype A sequences did not form a monophyletic clade in our tree, we removed them. We also removed a subtype A1 sequence that mixed with subtype A2. These steps produced a rooted tree of 1101 sequences which recovers all the subtypes as monophyletic. We used MD-Cat and the sampling years available from LANL to date the tree.

## 3 Results

### 3.1 Simulated angiosperms with hybrid rates

We first compare the methods in terms of the accuracy of the estimated divergence time of the most recent common ancestor (tMRCA). Our results show that MD-Cat outperforms wLogDate, RelTime, and BEAST-LN in all five rate scenarios (Fig. 2a). Both BEAST-LN and wLogDate severely overestimate the Angiosperm age for all scenarios. In scenario 1, where three BEAST rate priors are used, we observe that the strict and RLC models are better than the LN; however, they are still worse than wLogDate and far worse than MD-Cat. RelTime has a lower error than BEAST-LN and wLogDate, but, unlike all other methods, it underestimates the age. In contrast, MD-Cat has both low error (5.2% on average) and low bias: for scenario 1, it overestimates the age by 5 million years; for scenarios 2–5, it underestimates by 0.5, 1.6, 0.1, and 0.6 million years, respectively. Overall, MD-Cat is more accurate and less biased than the other methods in tMRCA estimation.

**Figure 2.**
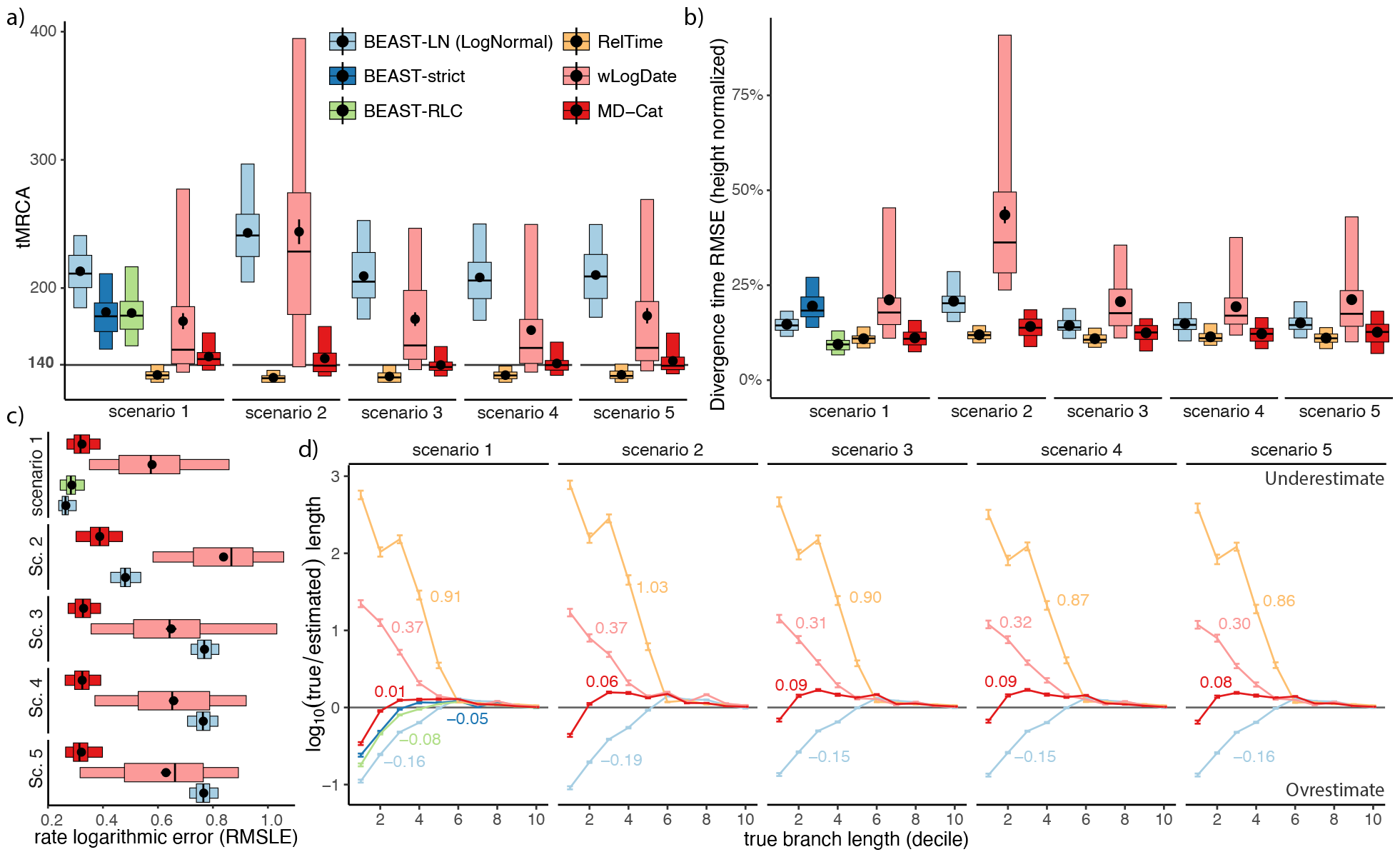
Comparison on the simulated Angiosperm dataset. The 5 scenarios differ in the ratio of substitution rates among some groups. Let *H* stand for herbaceous angiosperm, *W* for woody angiosperm, *A* for all angiosperm, *G* for gymnosperm. Scenario 1: ^*H*^/_*W*_ = 3. Scenario 2: ^*H*^/_*W*_ = 6. Scenario 3: ^*A*^/_*G*_ = 4 and ^*H*^/_*W*_ = 3. Scenario 4: ^*A*^/_*G*_ = 4 and ^*H*^/_*W*_ = 3 and *G* = *H*. Scenario 5: ^*A*^/_*G*_ = 4 and ^*H*^/_*W*_ = 3 and *G* = *W*. In scenario 1, three BEAST variants are tested. (a) The estimated tMRCA versus the true tMRCA = 140 (solid line). We show mean (black dots), and standard error (error bar), 25-75% percentiles (wide boxes), and 5-95% percentiles (narrow boxes) over 100 replicates. (b) The root mean squared error across all edges, normalized by tree height; settings as in (a) (c) Root-mean-squared base 10 logarithmic error (RMSLE) of the per-branch estimated substitution rates; statistics as in (a). Note that only methods that can estimate per-branch rates are included. (d) Bias in the estimate of branch lengths. We divide true branch lengths into deciles, and for each, show the mean and standard error of bias defined as the difference between log-transformed true and estimated lengths. The overall average is annotated beside each line. RelTime can return 0 length; these are replaced with 10^*−*5^ to allow log transformation (removing those results in a similar picture). Horizontal line at 0 indicates no bias.

**Figure 3.**
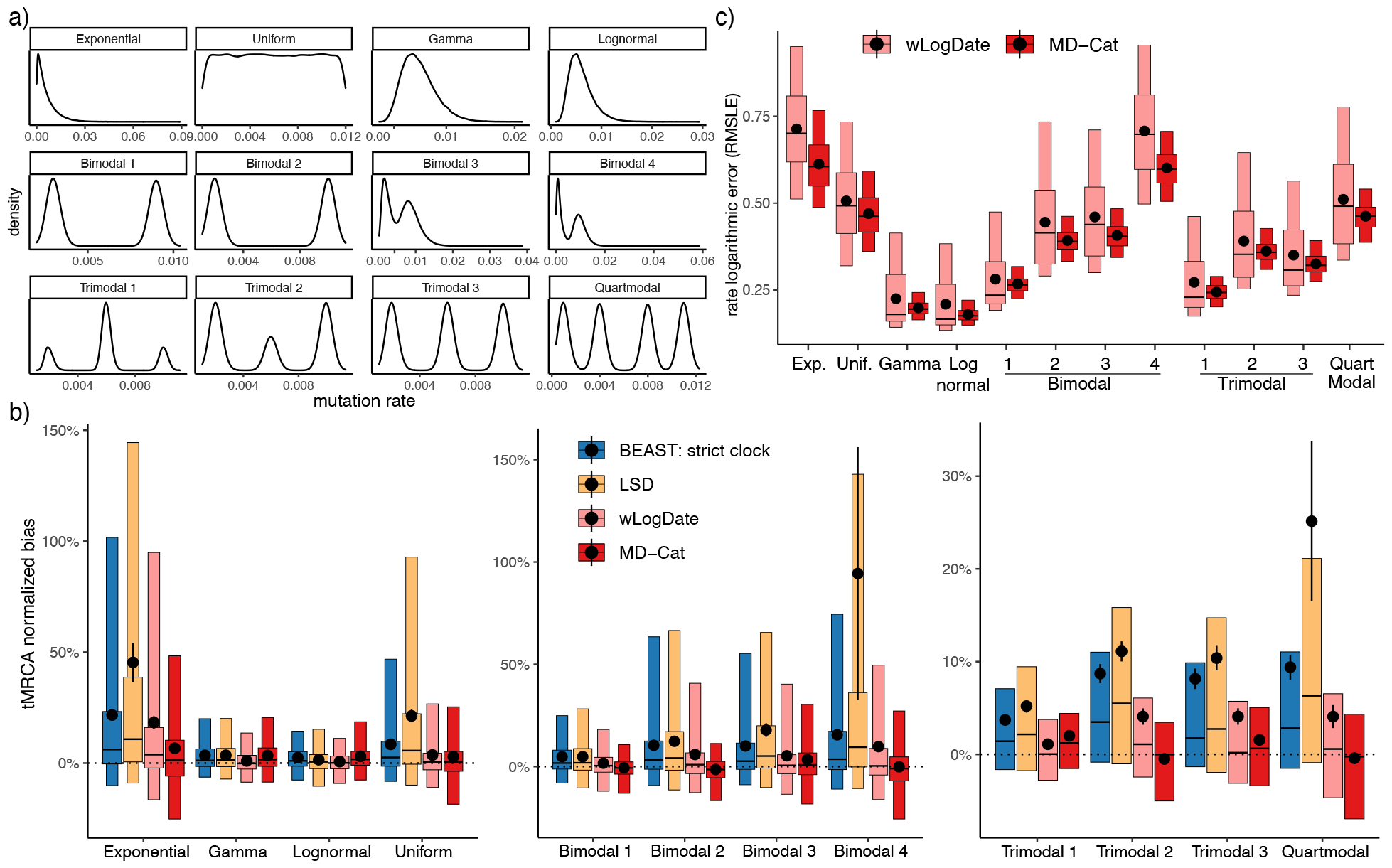
Comparison on the HIV simulated dataset. (a) The empirical distribution of the substitution rates simulated for each of the 12 clock models. (b) The bias in estimated root divergence time (tMRCA). The true tMRCA is fixed at0. Model conditions are divided into those with zero or one mode (left), two modes (middle), and three or more modes (right). Distributions are (same settings as Figure 2) over *n* = 400 data points (four tree conditions, each with 100 replicates). (c) Root-mean-squared logarithmic (base 10) error (RMSLE) of the per-branch estimated substitution rates of wLogDate and MD-Cat. BEAST is run with the strict clock model in all cases.

Next, we examine the mean divergence time error across all nodes using root-mean-squared-error (RMSE) divided by the tree height. Averaged over all five scenarios, RealTime has the least RSME (11.2%), closely followed by MD-CAT (12.5%) (Fig. 2b). The other methods have substantially higher errors, with the next best method, BEAST-LN (16%), substantially outperforming wLogDate (25.2%). Relative patterns of performance are similar across scenarios. In scenario 1, the BEAST-RLC prior outperforms other priors and slightly outperforms RelTime and MD-Cat (9.4% vs 10.9%, and 11.1%, resp.) BEAST-strict has the least accuracy among BEAST variants. The relative accuracy of BEAST priors matches our expectation given that local rates are used in this simulation.

In terms of the per-branch substitution rates, MD-Cat dramatically outperforms wLogDate (Fig. 2c). Comparison to BEAST depends on the scenario. BEAST-LN and BEAST-RLC both produce slightly more accurate per-branch rates compared to MD-CAT in scenario 1, which is the simplest scenario. As we move to more complex scenarios, the rates of BEAST-LN quickly become less accurate, leaving MD-CAT as the most accurate method. Note that unlike wLogDate and MD-CAT, which rely on input SU lengths computed by RAxML, BEAST co-estimates rates and SU lengths; thus, the relative accuracy of rates of BEAST does not necessarily match the relative accuracy of divergence time estimates.

MD-Cat also tends to estimate individual branch lengths better than other methods (Fig. S3). In particular, MD-Cat has lower (log-transformed) bias compared to other methods, especially for shorter branches (Figs. 2d). For short branches, wLogDate and RelTime underestimate the length, while all three BEAST variants and (to a lesser degree) MD-CAT overestimate. As branches get longer, the log bias reduces and eventually turns to underestimation for all methods. Across all branches, MD-CAT has the lowest log bias (e.g., 0.01 in scenario 1), followed by BEAST-strict (−0.05) and BEAST-RLC (−0.08). RelTime has a substantial tendency to underestimate the branch lengths, consistent with its underestimation of the tMRCA. BEAST variants overestimate for more branches than underestimate, leading to an overestimation of the tMRCA. Note that we measured the bias after a log transformation to ensure the error is not dominated by long branches. Measuring the bias without log transformation leads to different and less consistent patterns (Fig. S4). RelTime continues to have a high under-estimation bias (mean: 4 – 5 million years per branch). However, wLogDate becomes erratic, with the lowest bias in some scenarios and replicates and the largest bias in others. BEAST variants have relatively low levels of bias (*<* 1.2 million years on average), followed closely by MD-CAT (1.6 – 2.7 million years).

We next ask how the number of rate categories, *k*, impacts the results. Higher *k* values tend to usually (but not universally) increase accuracy, with the divergence time error ranging from 10-13% with *k ≥* 50 to 15-27 % with *k ≤* 10 (Fig. S5a). While our default value of *k* = 50 is among the best in accuracy, it is not universally the best choice as *k* = 100 often performs better. Regardless of *k*, the divergence time error of MD-CAT is lower than wLogDate and higher than RelTime (with one exception); however, comparison to BEAST-LN depended on *k*, with MD-Cat tending to be better for *k ≥* 25 and worse with *k ≤* 5.

We tested three methods to automatically select *k*: AIC, BIC, and cross-validation. Among these, crossvalidation has the lowest average error, followed by AIC, then BIC (Fig. S6a). Both AIC and BIC always select *k* in {10, 25, 50}, with *k* = 25 selected most often (Fig. S6b,c). Cross-validation selects a wider range of *k*, but tends to select *k ≥* 25 in the vast majority of cases (*k* = 100 in 42%, *k* = 50 in 26%, and *k* = 25 in 13% of replicates; see Fig. S6d). The accuracy of the *k* selected through either AIC, BIC, or cross-validation is slightly lower than simply using the default *k* = 50 (Fig. S6a); however, all of these three methods perform better than the average of all values of *k* other than 50. Using cross-validation to select *k* instead of the default results in 0.6% increase in the divergence time error across all 7,500 nodes of all replicates in all scenarios, providing an accurate (but slightly worse) alternative to the default *k* = 50.

### 3.2 Simulated HIV phylodynamics

All methods estimate the tMRCA most accurately on the two simple unimodal models (Gamma and Log-Normal), and substantially less accurately on the non-modal (Exponential and uniform) and multimodal models (Fig. 3b). The long-tailed exponential distribution and all the multimodal distributions except for Biomodal 1 and Trimodal 1 seem particularly challenging. The two multimodal distributions that are less challenging are symmetric around their mean and have less distance between modes. Distributions with long tails (compared to the mean) seem particularly challenging for all the methods.

Compared to the other methods, MD-Cat estimates tMRCA with comparable or better accuracy across all conditions (Fig. 3b). As expected, the strict-clock method, LSD, performs well on the unimodal cases (Gamma and LogNormal), but its accuracy drops dramatically on the other clock models. While wLogDate and BEAST-strict are more robust than LSD, they both overestimate the tMRCA across all clock models (the average bias is 5.0% and 8.9% for wLogDate and BEAST, respectively, averaged over all models) and especially for the nonmodal and multi-modal distributions. For example, while the biases are less than 2.5% for both methods under the LogNormal distribution, under the challenging Binomial 4 model, the mean bias is 15.5% and 9.8% for BEAST and wLogDate, respectively. MD-Cat overestimates the tMRCA in seven models, among which only Exponential has a somewhat high bias (6.7%), and the rest (Gamma, LogNormal, Uniform, Bimodal 3, Trimodal 1, and Trimodal 3) have a bias below 3.5%. In the other 5 conditions, MD-Cat slightly underestimates tMRCA, with the average bias at 1.4% for Bimodal 2 and *<* 1% in other cases (Bimodals 1 and 4, Trimodal 2, and Quartmodal). Averaged across all models, MD-Cat has a small overestimation bias of 1.7%. MD-Cat also has fewer cases of extreme error; e.g., MD-CAT has tMRCA errors above 50% in 50 replicates across all conditions compared to 213 and 137 replicates with BEAST and wLogDate. Finally, the advantage of MD-Cat over other methods is most clear for the inter-host tree models, which tend to be more challenging (Figure S8a).

The error of divergence time estimation shows similar patterns (Fig. S8b). In the two unimodal models, all methods are relatively accurate and have similar accuracy. MD-Cat is more accurate than the other methods for Exponential (non-modal) and multi-modal clock models. The average improvements of MD-Cat over the second-best method range from small (e.g., 0.3% for Trimodal 1 model) to substantial (3% or more for Exponential, Bimodal 2, and Bimodal 4; see Fig. S8c). Across all clock models, the average errors of LSD, wLogDate, BEAST, and MD-Cat are 15%, 8.0%, 8.1%, and 6.7%, respectively. MD-CAT also has fewer cases of very high error. For example, the 95-percentile error of LSD, BEAST, and wLogDate for the Exponential model are 70%, 48%, and 50%, resp., compared to 22% for MD-Cat. Comparing the estimated substitution rates by wLogDate and MD-Cat confirms the same relative performance of the two methods, where MD-Cat has a lower average error in all clock models (Fig. 3c).

The choice of *k* has only a moderate impact on the MD-Cat divergence time accuracy (Fig S5b). As long as *k ≥* 5 is chosen, little change in accuracy is observed. Going from *k* = 2 to *k* = 5 results in a significant improvement (mean RMSE goes from 0.065 to 0.055, and *p*-value= 0.00003 according to a paired t-test); in contrast, changing *k* = 5 to *k* = 10 or *k* = 50 *increases* mean error but not significantly (*p* = 0.59, *p* = 0.13). Thus, while default *k* = 50 is not the best choice, it is close (mean RMSE 0.057). Depending on the condition, the best *k* can vary from as low as 5 to as high as 100 (Fig S5b). The three methods to select *k* (AIC, BIC, and cross-validation) have similar accuracy on this dataset, and their relative performance depends on the tree model (Fig. S7a). Both AIC and BIC tend to select small values for *k* (e.g. 2,5, or 10), with one exception where AIC selects *k* = 25 (Fig. S7b,c). In contrast, cross-validation can pick a wide range of *k* values (Fig. S7d). Overall, AIC is the best method, followed by cross-validation, then BIC, but the difference is statistically insignificant (p=0.89 for AIC versus cross-validation and p=0.9 for BIC versus cross-validation). Using one of these methods to select *k* is often (but not always) slightly *better* than the default *k* = 50 and other choices of *k* (Fig. S7a).

On this dataset, we also examined the accuracy of the confidence intervals (CI) estimated by MD-CAT, focusing on the challenging trimodal rate distributions (Fig. S10). As expected, because the accuracy of the point estimate is lower for inter-host compared to intra-host models (Fig. S8c), the CIs of both branch lengths and divergence times are wider on the inter-host models (Fig. S10a). Nevertheless, these trimodal distributions of the rates present challenging cases for CI estimation. Overall, 20% of branches and 22% of node divergence times fall outside of the 95% CI (Fig. S10b), showing CIs are too narrow. Nevertheless, even when the CI does not strictly include the true value, in about 80% of cases, the true value would have been inside if the CI was wider by around ^1^/_3_ (Fig. S9c). Thus, when CI fails to include the true value, in most cases, it is not far from capturing it.

Comparing the running times, LSD is extremely fast, finishing each replicate in less than a second. Among the other methods (Fig. S9 in Supplementary), wLogDate is the fastest method with an average running time of 15 minutes, followed by MD-Cat at 30 minutes, which is more than 5 times faster than BEAST at 159 minutes. Note that the topology is fixed in all cases. There is no substantial difference in running time among different clock models. Runtime for bootstrap by MD-Cat is unsubstantial, adding only 2 minutes to the overall runtime to obtain 1000 bootstrap replicates.

### 3.3 Dating of the real angiosperm data

Beaulieu et al. (2015) showed that BEAST with the LogNormal clock model estimates a very old age for the crown angiosperm (median 232 Mya; 95% HPD 210–256), and this estimate is much older than evidence from fossils. In particular, the earliest available fossils reliably identified as ancestral angiosperms are dated to 140 mya. The 100 million years gap between the BEAST estimate and the earliest known fossil led to the suspicion that BEAST could be overestimating dates due to the use of an incorrect clock model. The authors concluded that sources of bias in BEAST mean that we cannot rule out a much more recent age, closer to 140 mya, than what BEAST estimates. MD-Cat estimates the divergence time of angiosperms to be 189 Mya (Fig. 4). Although this estimated age is still substantially older than the earliest available fossil, the gap is reduced by half relative to BEAST. Moreover, our estimate is within the range of previous estimates by molecular dating, which ranged between 175 and 251 Mya (Beaulieu et al., 2015; Clarke et al., 2011; Foster et al., 2017; Zeng et al., 2014), and is on the younger side. The estimated branch rates form a bimodal (or perhaps trimodal) distribution (Fig. 4) that would not fit commonly used unimodal distributions such as LogNormal. This distribution has a substantial number of branches with much higher rates (close to 0.0025) than typical branches, which have rates distributed around 0.00055. Some of these high-rate branches are close to the root of angiosperms; e.g., the stem branch leading to the sister clade of angiosperms has a high rate. Among herbaceous angiosperms singled out by Beaulieu et al. (2015) as potential sources of increased rate (Nymphaeales, Piperales, Monocotyledonae, and Ceratophyllum), some but not all branches have clearly elevated rates; e.g., branches leading to *Trithuria* at the base of Nymphaeales, *Piper* among Piperales, and the domesticated *Oryza* among monocots have elevated rates. Most strikingly, the MRCA of all monocots and the MRCA of all non-Araceae monocots both have rates close to 0.0025, which is 5*×* the typical rate across the tree. These branches, close to the base of angiosperms, likely have a major impact on its estimated tMRCA; lowering their rate would put the angiosperm tMRCA further back in time.

**Figure 4.**
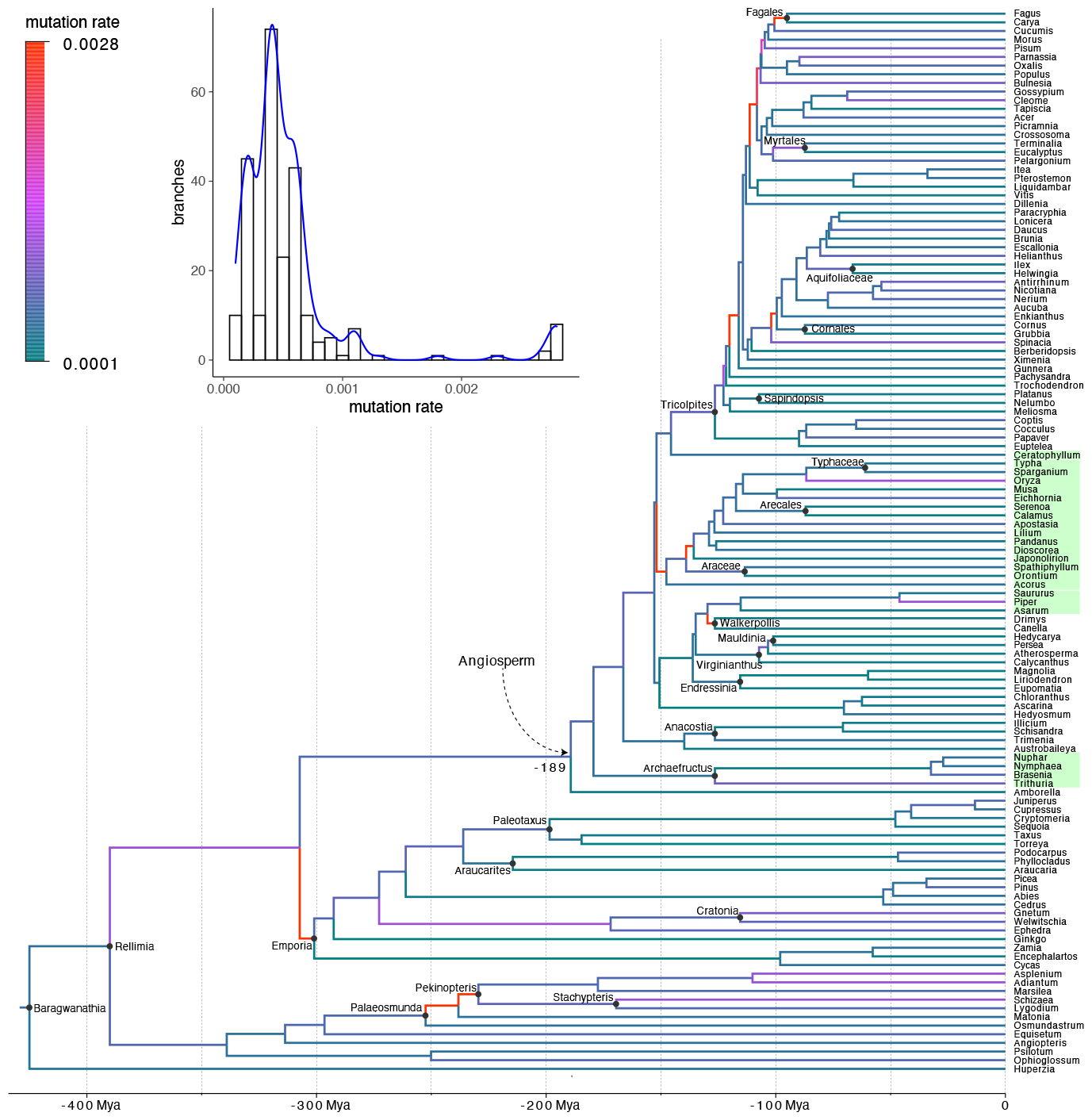
Dating land plants using MD-Cat. Branches are colored by the corresponding estimated substitution rate, with the histogram and density plot of the rates shown as inset. Calibration points are shown with solid dots and the corresponding name. The age of angiosperm is estimated at 189 Mya. Four herbaceous groups singled out by Beaulieu et al. (2015) (Nymphaeales, Piperales, Monocotyledonae, and Ceratophyllum) are highlighted in green.

### 3.4 Dating of the real combined HIV subtypes

Our estimate of the tMRCA of the entire HIV-1 M group is 1922, consistent with the hypothesis about the Kinshasa spread of HIV-1 in 1920s proposed by many analyses (Faria et al., 2014; Junqueira and de Matos Almeida, 2016; Keele et al., 2006; Korber et al., 2000). Also, according to these earlier studies, the subtype B lineage originated in Kinshasa during 1940s but was detected outside Africa only in 1960s (Faria et al., 2014; Junqueira and de Matos Almeida, 2016). Molecular dating methods often place the tMRCA for subtype B in between these two events, i.e., in 1950s (Abecasis et al., 2009; Korber et al., 2000; Patiño-Galindo and González-Candelas, 2017; Wertheim et al., 2012). Consistent with these results, MD-Cat estimates the tMRCA of subtype B to be 1950, which is close to the proposed true origin in 1940s. In addition, our estimates of the tMRCA of most of the other subtypes are also within the confidence interval of previous works (Fig. 5a and Table 1). The average substitution rate across all subtypes is estimated at 0.0032, and we observed no substantial difference among the subtypes (Fig. 5b). Tracking the substitution rates through time (Fig. 5c) by averaging all branches spanning each year showed consistent estimates across years until the early 2000s when the rate started to drop, a point that we will return to in the discussion.

**Table 1:**
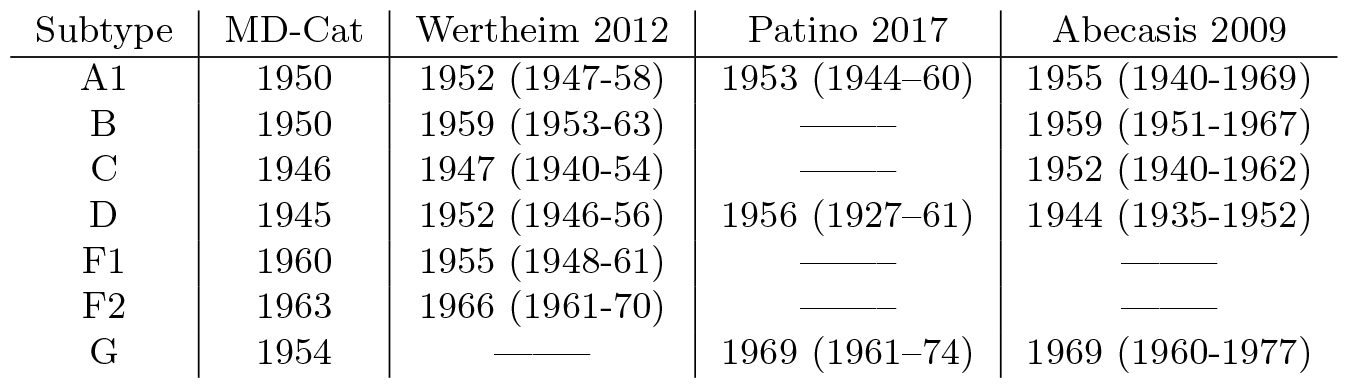
The estimated divergence time of the HIV-1 subtypes.

**Figure 5.**
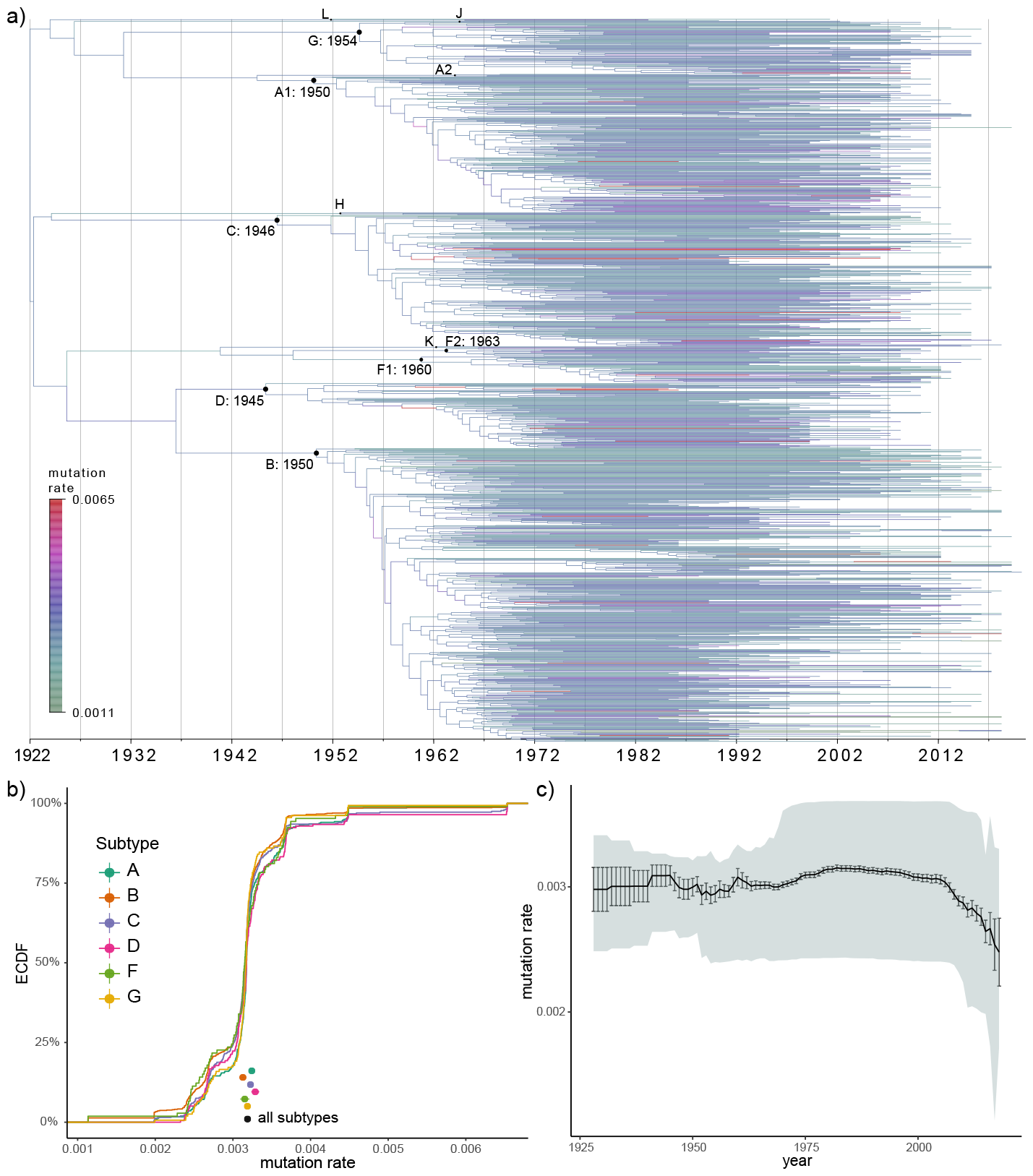
Dated HIV tree. a) The chronogram estimated by MD-Cat. Branches are colored by the estimated substitution rate. b) The distribution of rates separated for the major subtypes, shown as an empirical cumulative distribution (ECDF). The mean and standard error of each subtype and all subtypes combined are also shown as dots and error bars (with arbitrary positions along the y-axis); only branches below the MRCA of each subtype are included. c) Rates across time. For each year, we computed the distribution of rates of all branches that span that entire year; we show the mean and standard error as a line and error bars and 10–90% quantiles as shaded area.

## 4. Discussion

We introduced a phylogenetic dating method using the flexible categorical model. We formulate the dating problem as maximum likelihood inference and solve it using an EM-based algorithm. Although the likelihood function is non-convex, the objective function in the M-step is convex on *ω* and *τ* separately. We use block coordinate descent to alternatively optimize *ω* and *τ*, and show that each iteration can be solved efficiently. Together, these strategies greatly reduce the running time of the MD-Cat method.

### 4.1 Further observations on the data

On both simulated datasets, MD-Cat outperformed other methods in most clock models according to most metrics, especially tMRCA time. The improvements were most visible in the model conditions with complex clock models. Comparing the accuracy of MD-Cat and wLogDate, we observed the most dramatic improvements under the hardest model conditions where either there was a sudden extreme rate shift (angiosperms) or the overall rate distribution had high variance (HIV). Interestingly, the non-modal long-tailed Exponential distribution was one of the most challenging conditions for all methods and also experienced one of the largest improvements. Somewhat surprisingly, the uniform distribution was also relatively difficult for LSD and BEAST (strict clock). Not all bimodal distributions were equally challenging. Bimodal 1 with two completely separate peaks but low variance around peaks was easy while the bimodal 4 condition with a higher variance around the peaks was challenging. Overall, all methods had more difficulty dealing with long tails of high rates, and MD-Cat dealt with this difficulty better than other methods. Thus, the benefit of using the categorical model seemed related to its ability to capture outlier branches that exhibit high rates due to biological change or artefactual reasons (e.g., model misspecification or missing data). Interestingly, tails of outlier branches with elevated rates were observed on both biological datasets.

On the angiosperm data, the tMRCA estimated by MD-CAT (189 mya) was younger than most other reported ages but still older than the age of the youngest fossil (140 mya). Consistent with a younger age, MD-Cat found several branches with elevated rates close to the base of angiosperms, including a tail of some branches with very high rates. The remaining gap in the tMRCA age could be due to inaccuracies in the MD-Cat estimated date. Perhaps MD-Cat underestimates the rates for several branches close to early angiosperm diversification. However, with the given fossil dates (treated as fixed dates) getting to 140 mya requires very high rates after the angiosperm sperm diversification *and* a lowered rate on the branch above it. We note that in the angiosperm *simulations* where the tMRCA is set to 140 mya, MD-Cat never estimates an age as old as 189 mya. This is in contrast to BEAST, which almost always estimated the tMRCA at 180 mya and often older than 200 mya. Thus, our results indicate that an older age for angiosperm (somewhere between 140 and 189 mya) than implied by the fossil record cannot be rejected; the available fossils may belong to species that lived subsequent to the early angiosperm divergences.

On the HIV real data, MD-Cat was generally in agreement with prior publications. We saw little difference between the substitution rates of subtypes, but all subtypes included outlier branches with much higher rates (up to 0.006). Similarly, the rate of substitutions seemed mostly consistent through time, but there was a drop in rates circa 2000. This period coincides with the wider availability of HIV antiviral drugs, which can stop the evolution of the virus while taken by an individual. Thus, the treatments may lead to lowered observed substitution rates at the population level by pausing the accumulation of substitutions, which resumes when individuals stop taking the treatment, which happens often. However, this trend can also be an artifact of the lower number of branches spanning recent years (and hence the higher variance in recent years). Nevertheless, these potential connections to epidemiological factors provide an interesting insight. Further studies should examine if the rate reduction in recent years holds up if the sampling of recent years is further improved.

### 4.2 CAT models and the choice of *k*

Our method allows multi-modal uncorrelated substitution rate distributions using categorical models. One approach to allow multi-modality is to use local clocks. For example, Fourment and Darling (2018) use independent relaxed clocks (e.g., LogNormal distribution) for different clades of the tree defined *a priori*. Much earlier, Drummond and Suchard (2010) had proposed the random local clock model (which we tested) that eliminated the need to prespecify the clades. They used a Bayesian selection method to sample over many possible random local clocks, with different clade boundaries. Regardless of whether clades of shared rates are prespecified, these local models allow multi-modality using a very different approach than ours because the rates are clade-dependent. In contrast, we assume i.i.d rates but allow multi-modal distributions using categorical approximation. Our use of categorical models for i.i.d rate variation is similar to the model that Höhna et al. (2019) have proposed for the problem of diversification rate heterogeneity across the tree. Categorical models (sometimes abbreviated to CAT) have been used extensively in various phylogenetic modeling tasks. Foster (2004) developed a model where branches of the tree draw the base frequency parameters of the sequence evolution model from a set of vectors. Instead of heterogeneity across the tree, Lartillot and Philippe (2004) assumed sites of the alignment are distributed according to a mixture of distinct classes, each given its own substitution matrix. Later, Blanquart and Lartillot (2008) extended this approach to allow base frequency heterogeneity across both sites and branches. Most relevant to our work, Heath et al. (2012) adopted rate categories for joint inference of the tree and its divergence times. Under their model, branch rates are drawn from a prior distribution consisting of clusters of rates, each cluster with its own distribution. These methods use Bayesian MCMC to explore the posterior space of the specified models, including the assignment of branches or sites to categories. Since MCMC sampling does not require optimization, the models used in these methods tend to be more complex than ours (e.g., *k* is often a random variable). However, MCMC with many parameters can lead to slow convergence, a limitation that our ML-based approach can ameliorate.

Likelihood-based approaches also exist. The discrete clock model of Fourment and Holmes (2014) (FH14) uses categories to allow varying substitution rates across the tree. In this model, each branch is independently assigned a rate from a set of possible rates. Despite some similarities, the FH14 and our model have a major difference. The FH14 discrete clock model assigns each branch to one of the *k* rate categories. In contrast, we assume each branch draws its rates from a categorical rate distribution. Thus, in the FH14 model, each branch has a single rate assigned at the end, whereas, in our approach, we get a probability for each branch belonging to each rate category. This simple difference has a major practical implication. In the hard-assignment model of FH14, matching rates to categories becomes a combinatorial problem, which authors heuristically resolve using a genetic algorithm. The soft assignment formulation of our approach gives us a continuous optimization problem, which we show can be solved efficiently using the EM algorithm.

Methods based on the CAT model have proposed several approaches for selecting the number of categories. Lartillot and Philippe (2004) used the Dirichlet process to sidestep the need to predefine the number of categories, effectively allowing a countably infinite *k*. Thus, the space of all possible mixtures is explored in a Bayesian fashion, guided by a prior distribution and hyperparameters that influence heterogeneity. This approach has been since adopted by many other Bayesian methods based on the CAT model (Heath et al., 2012; Huelsenbeck et al., 2006). In contrast, Fourment and Holmes (2014) used traditional model selection techniques, such as the Akaike Information Criterion (AIC) to find the optimal *k*, an approach that we have also implemented inside MD-CAT.

MD-CAT uses a large fixed *k* by default, and this strategy worked well empirically in our simulations. A large *k* value is justified because we use categorical models to approximate a continuous distribution. It is worth considering what could happen if the true evolutionary process had *k*^*∗*^≠ *k* true rate categories. If *k*^*∗*^ *> k*, we will have an approximation error. We have made this unlikely by selecting *k* = 50 and encourage users to use even higher *k* values for much larger trees than explored here. In contrast, when *k*^*∗*^ *< k*, our method can select *ω* values that are extremely similar. If the *ω*s found by MD-CAT include *m ≤ k* values in the [*r, r*(1 + *ϵ*)] range (for a small *ϵ*), the number of distinct rates is effectively reduced to *k − m* + 1, with the probability of the rate being (approximately) equal to *r* set to ^*m*^/_*k*_. Thus, overall, our recommendation is to fix *k* to a large value (which clearly should not exceed the number of edges in the tree). We note that having a large number of rate categories does not seem to cause a problem in terms of accuracy (Fig. S5) but can impact the running time. However, using a large *k* is still far more scalable than a model selection technique such as AIC, BIC, or cross-validation.

### 4.3 Limitations and future work

Despite its improved empirical accuracy, many theoretical and computational questions related to MD-Cat remain unanswered. Foremost, MD-Cat can suffer from over-parameterization. As noted before, time and rate are inseparable from molecular data, even with the aiding information from calibration points. In general, approximating a continuous distribution requires a large number of rate categories (*k*). However, the data size is small compared to the number of unknown parameters (2*n −* 2 branch length observations and *O*(*n*) calibration points versus 2*n −* 2 + *k* parameters). Therefore, the parameters are unidentifiable in general, pointing to the difficulty of getting the correct age for every node. MD-CAT pre-estimates the mean substitution rate and uses a two-round optimization strategy to alleviate this challenge. Nevertheless, our evidence that the inference of a substantial number of distinct rates is possible despite the large space is mostly empirical. We leave the theoretical analysis of identifiability of *ω* and *τ* under this categorical model and different combinations of calibration points for future work.

In our formulation, we incorporated internal node calibration data as point values. Since fossil records do not always correspond to the exact internal node available, two types of improvements can be incorporated in the future. First, the equality constraints can be changed to inequality, a change that would be relatively easy to incorporate. But going further, many of the modern dating methods allow the age of the internal nodes to be specified as a distribution, allowing for uncertainty. Such a framework can also be incorporated into the likelihood framework described here by adding an internal node age likelihood term. If these likelihood terms are considered independent, the formulation becomes conceptually simple, leaving only the challenge of making sure the optimizer is still able to converge.

Beyond the model and the constraints, the inference procedure could also improve. For scalability purposes, we use the inferred tree in substitutions per site as input and use a simple Gaussian model for branch length estimation error. However, a more direct method is to incorporate the categorical clock model into the tree inference from sequence data and maximize the joint likelihood function. While this approach avoids assuming a Gaussian model for branch length estimation error, we note that it may exacerbate the over-parameterization problem, as all the parameters (tree topology, GTR parameters, per-site, and per-branch rate heterogeneity) must be co-estimated. Future works should explore this approach, both theoretically and empirically. Finally, rooting phylogenies is a non-trivial problem and is also related to the problem of clock model selection. The current formulation of MD-Cat only works on rooted phylogenetic trees, but the generalization of MD-Cat to unrooted trees is a straight-forward: solve the optimization problem for each possible rooting and select the root position that has the maximum likelihood. Such an approach should be explored in future works, together with an updated formulation for MD-Cat to add a parameter that determines the optimal root placement on each branch.

Our study can also be extended in the future. First, to facilitate the comparison between different methods, we used the true topology with estimated branch lengths and left the study of the impact of incorrect topology on the dating methods for future works. In addition, testing under more complex clock models, such as those that allow continuous rate change, is also a topic for future work.

## 5. Acknowledgement

This work was supported by the National Institutes of Health (1R35GM142725).

