## Supplementary for "Expectation-Maximization enables Phylogenetic Dating under a Categorical Rate Model"

### Supplementary information of MD-CAT

**B Supplementary figures and tables** **7**

#### List of Algorithms

1 The Lagrange method to solve a subproblem $\mathcal{P}^{(i)}$ defined by the active set $\mathcal{A}^{(i)}$ . . . 4
2 The active-set method to solve the problem $\mathcal{P}$ defined in Eq. S1 . . . 6

#### List of Tables

S1 Parameters and statistics of the 12 clock models . . . 7
S2 The 5 scenarios of the simulated angiosperms . . . 8
S3 The estimated divergence time of the HIV-1 subtypes . . . 8

#### List of Figures

S1 Log-likelihood versus iterations of the EM algorithm on the first replicate using 200 initial points. The thick line shows the best final log-likelihood. . . . 9
S2 Effective sample size (ESS) of the posterior for root age in the 5 different angiosperm tree scenarios when analyzed on BEAST. Different values are shown when analysis was run for 10, 20, and 30 million generations. ESS above the yellow horizontal line (y = 100) report the minimum required to demonstrate MCMC convergence, while those above the green line (y = 200) are more clearly converged. . . . 10
S3 Estimated versus true branch length. Each dot shows the mean and standard error over all 100 replicates for that branch in the true tree. Both axis are in log scale. . . . 11
S4 The distribution of the mean difference between true and estimated branch length across all branches of each replicate, indicating bias. We show mean (black dots), and standard error, 25-75% percentiles (wide boxes), and 5-95% percentiles (narrow boxes) over 100 replicates. . 12
S5 Analysis of the sensitivity of MD-Cat to the number of rate categories ($k$) for (a) the Angiosperms simulated data, and (b) the HIV simulated dataset under the three trimodal clock models. The default $k$ used in all other analyses is 50. We compare methods in terms of the divergence time error, defined as the root mean squared error of divergence times of all nodes versus true divergence, normalized by tree height. . . . 13

S6 Selection on $k$ for the Angiosperms simulated data using AIC, BIC, and cross-validation. (a): The divergence time error (absolute difference of true and estimated normalized by tree height) across all branches. b,c,d: The selected $k$ distribution across all runs for AIC, BIC, and cross-validation, respectively. . . . . . 14
S7 Selection on $k$ for the HIV simulated data using AIC, BIC, and cross-validation. (a): The divergence time error (absolute difference of true and estimated normalized by tree height) across all branches. b,c,d: The selected $k$ distribution across all runs for AIC, BIC, and cross-validation, respectively. . . . . . 15
S8 Detailed comparisons on the HIV simulated dataset. a) Normalized tMRCA error, broken down by tree condition and clock model. Since LSD can have extremely high errors, we removed it from the figure to help readability. Narrow boxes show 5-95 percentiles and wide boxes show 25-75 percentiles. Bar shows median. Mean and standard error are also shown as dots and error bars. b) Divergence time RMSE (normalized by tree height). Mean/se omitted due to high mean error for LSD. Gamma and Lognormal are unimodal. Uniform and Exponential are nonmodal. c) Same as (b) with tree models and clock models separated. . . 16
S9 Comparison of the running time of BEAST, MD-Cat, and wLogDate separated into four different clock model groups. The y-axis represents the time in minutes and is shown in the log-scale. All analyses were run on Intel Xeon CPU E5-2670 (103,597,240 kB of memory). MD-Cat runtime does not include CI calculation and SU branch length estimation but those would add a negligible amount: about 2 minutes for CI estimation and several seconds for SU branch length estimation using RAxML. . . . . . 17
S10 Evaluating support values. a) Confidence interval (CI) estimated by MD-Cat for branch lengths (Left) and divergence times (Right). Blue: CI includes the true value; red: it does not. b) The portion of branches (Left) or nodes (Right) that fall outside the 95% CI, shown separately for 10 deciles of true branch length / divergence time. c) For nodes where the true value falls out of CI, we show how far it falls as a percentage. We show the distribution of $|t - c|/t$ where $c$ is the CI boundary which fails to capture the true value and $t$ is the true divergence time. . . . . . 18

### A Supplementary text

#### A.1 Solving for $\omega$ in the M-step

Here we show the algorithm to solve $\omega$ in the first round of MD-Cat where we add a constraint on $\omega$ so that its average is a constant $\mu$. We note that the optimization problem in the second round is simpler and can be solved using a similar algorithm.

##### A.1.1 The optimization problem

Recall that we use block coordinate descent to solve the M-step of MD-Cat. In each iteration, if we let $\tau$ fixed, then the optimization problem is reduced into the following:

$$\mathcal{P} : \min_{\omega} \sum_{i=1}^{N} \sum_{j=1}^{k} \frac{q_{ij}}{\hat{b_i}} (\hat{b_i} - \omega_j \tau_i)^2 \tag{S1}$$

such that $\omega \geq \epsilon$ and $\sum_{j=1}^{k} \omega = k\mu$, where $\epsilon$ and $\mu$ are constants and $0 \leq \epsilon < \mu$. It is trivial to see that the optimization problem $\mathcal{P}$ is convex. Therefore, we can find the global optimal $\omega^*$ of $\mathcal{P}$ using the active-set method [1] as detailed below. In addition, thanks to the simple weighted least-square form of the objective function, below we show that each iteration of our active-set algorithm can be solved in linear time, which is $O(N + k)$.

##### A.1.2 Overview of the active-set method

Recall that a *feasible point* of an optimization problem is a point that satisfies all the problem's constraints. If $\omega^{(i)}$ is a feasible point of $\mathcal{P}$, then its *active-set* $\mathcal{A}^{(i)}$ is defined as:

$$\mathcal{A}^{(i)} = \{j | \omega_j^{(i)} = \epsilon\} \tag{S2}$$

Starting with a feasible point $\omega^{(1)}$ and its active-set $\mathcal{A}^{(1)}$, the active-set algorithm repeat the following procedure in each iteration (i) until the optimal point is found:

- Solve the equality-constraint problem $\mathcal{P}^{(i)}$ defined by the active-set $\mathcal{A}^{(i)}$ (the formal definition of $\mathcal{P}^{(i)}$ will be shown later).
- Use $\omega^{(i)}$ and the optimal point $\omega^{*(i)}$ of $\mathcal{P}^{(i)}$ to find a new *feasible* point $\omega^{(i+1)}$ that is closer to the optimal point of $\mathcal{P}$ than $\omega^{(i)}$.
- Compute the active-set $\mathcal{A}^{(i+1)}$ of $\omega^{(i+1)}$.
- Replace $\omega^{(i)}$ with $\omega^{(i+1)}$ and $\mathcal{A}^{(i)}$ with $\mathcal{A}^{(i+1)}$, then repeat the procedure.

##### A.1.3 The subproblem $\mathcal{P}^{(i)}$ and the Lagrange method

In iteration $(i)$ of the active-set method, we define and solve the following equality-constraint optimization problem:

$$\mathcal{P}^{(i)} : \min_{\omega} \sum_{i=1}^{N} \sum_{j=1}^{k} \frac{q_{ij}}{\hat{b_i}} (\hat{b_i} - \omega_j \tau_i)^2 \tag{S3}$$

such that $\omega_j = \epsilon, \forall j \in \mathcal{A}^{(i)}$ and $\sum_{j=1}^{k} \omega = k\mu$.

$\mathcal{P}^{(i)}$ can be solved *analytically* by introducing Lagrange multipliers:

$$L = \sum_{i=1}^{N} \sum_{j=1}^{k} \frac{q_{ij}}{\hat{b_i}} (\hat{b_i} - \omega_j \tau_i)^2 - \eta \left( \sum_{j=1}^{k} \omega_j - k\mu \right) - \sum_{j \in \mathcal{A}^{(i)}} \lambda_j (\omega_j - \epsilon), \tag{S4}$$

**Algorithm 1** The Lagrange method to solve a subproblem $\mathcal{P}^{(i)}$ defined by the active set $\mathcal{A}^{(i)}$

```
function SolveLagrange(A^(i))
    ω ← []                                    ▷ initialize to an empty list
    λ ← {}                                    ▷ initialize to an empty dictionary
    compute η using Eq. S7
    for j ∈ [k] do
        if j ∈ A^(i) then
            compute λ_j using Eq. S8
            λ[j] ← λ_j
            ω_j ← ϵ
        else:
            compute ω_j using Eq. S6
        append ω_j to ω
    return ω, η, λ
```

where $\eta$ is the Lagrange multiplier of the constraint $\sum_{j=1}^{k} \omega = k\mu$ and each $\lambda_j$ is the Lagrange multiplier of the constraint $\omega_j = \epsilon$. We will use $\lambda$ to represent the vector containing all these Lagrange multipliers. Because $\mathcal{P}^{(i)}$ is convex, $L$ has a unique critical point which also gives the unique solution to $\mathcal{P}^{(i)}$. We have:

$$\frac{\partial L}{\partial \omega_j} = \sum_{i=1}^{N} \left( \frac{2q_{ij}\tau_i}{\hat{b_i}} (\omega_j \tau_i - \hat{b_i}) \right) - \eta - \lambda_j \mathrm{I}_{j \in \mathcal{A}^{(i)}}, \tag{S5}$$

where I is the indicator function. We can find the critical point of $L$ by setting its partial derivatives with respect to each $\omega_j$ and $\lambda_j$ to 0. Equivalently, we set $\frac{\partial L}{\partial \omega_j}$ to 0 and use the original constraints $\omega_j = \epsilon, \forall j \in \mathcal{A}^{(i)}$ and $\sum_{j=1}^{k} \omega = k\mu$ to form a system of equations on $\omega$ and $\lambda$. Below we show how to solve this system of equations.

Let $\omega^*$, $\eta^*$, and $\lambda^*$ denote the critical point of $L$. If $j \in \mathcal{A}^{(i)}$ then $\omega_j^* = \epsilon$. Otherwise, $\omega_j^*$ can be solved by setting $\frac{\partial L}{\partial \omega_j}$ to 0. Thus, we have

$$\omega_j^* = \begin{cases} \epsilon & j \in \mathcal{A}^{(i)} \\ \frac{a_j}{c_j} + \frac{\eta^*}{k c_j} & j \notin \mathcal{A}^{(i)}, \end{cases} \tag{S6}$$

where $a_j = 2\sum_{i=1}^{N} q_{ij}\tau_i$ and $c_j = 2\sum_{i=1}^{N} \frac{q_{ij}\tau_i^2}{\hat{b_i}}$. Substitute this equation to the constraint $\sum_{j=1}^{k} \omega_j^* = k\mu$ and solve for $\eta^*$, we have:

$$\eta^* = \frac{k^2\mu - k\sum_{j=1}^{k} \frac{a_j}{c_j} - k\epsilon|\mathcal{A}^{(i)}|}{\sum_{j=1}^{k} \frac{1}{c_j}}, \tag{S7}$$

where $|\mathcal{A}^{(i)}|$ denotes the cardinality of $\mathcal{A}^{(i)}$. Next, substitute $\omega_j^* = \epsilon$ to Eq. S5 and set $\frac{\partial L}{\partial \omega_j}$ to 0, we can solve for each $\lambda_j^*$ where $j \in \mathcal{A}^{(i)}$:

$$\lambda_j^* = c_j\epsilon - \frac{\eta^*}{k} - a_j \tag{S8}$$

Substitute $\eta^*$ in Eq. S7 to Eq. S8 and Eq. S6, we obtain the analytical solution to all $\lambda_j^*$ and $\omega_j^*$. Obviously, this analytical solution can be computed in $O(N+k)$, as shown in Algorithm 1.

##### A.1.4 Computing $\omega^{(i+1)}$ and $\mathcal{A}^{(i+1)}$

Let $\omega^{(i)}$ and $\mathcal{A}^{(i)}$ denote the feasible point and its active-set found in iteration (i) of the active-set algorithm and let $\omega^{*(i)}$ denote the optimal point of $\mathcal{P}^{(i)}$, $\eta^*$ and $\lambda^{*(i)}$ denote its optimal Lagrange multipliers. Depending on the characteristics of $\omega^{*(i)}$ and $\lambda^{*(i)}$, we can find $\omega^{(i)}$ and $\mathcal{A}^{(i)}$ as follows.

**Case 1: $\omega^{*(i)}$ is feasible to $\mathcal{P}$ and $\lambda^{*(i)} \geq 0$** In this case, $\omega^{*(i)}$ is also the optimal point of $\mathcal{P}$, according to the KKT condition. We simple return $\omega^{*(i)}$.

**Case 2: $\omega^{*(i)}$ is feasible to $\mathcal{P}$ and $\exists\lambda_j^{*(i)} < 0$** In this case, set $\omega^{*(i+1)}$ to $\omega^{*(i)}$ and find the constraint $j$ that has the most negative $\lambda_j^{*(i)}$ and remove it from $\mathcal{A}^{(i)}$ to get $\mathcal{A}^{(i+1)}$ (i.e. relax the "useless" constraint).

**Case 3: $\omega^{*(i)}$ is infeasible to $\mathcal{P}$** In this case, we search for a new feasible point $\omega^{(i+1)}$ that is closer to the optimum and update the active-set. To this purpose, we start from the previous feasible point $\omega^{(i)}$ and move it as close as possible to $\omega^{*(i)}$ on the direction $d = \omega^{*(i)} - \omega^{(i)}$ such that the new point is still feasible. In other words, we need to find the largest number $\alpha \in [0, 1]$ such that

$$\omega_j^{(i)} + \alpha(\omega_j^{*(i)} - \omega_j^{(i)}) \geq \epsilon, \forall j \in \mathcal{A}^{(i)} \tag{S9}$$

Let $V = \{j|\omega_j^{*(i)} < \epsilon\}$ denote the *violated set* of $\omega^{*(i)}$. Because $\omega^{(i)}$ is feasible, it is easy to see that Eq. S9 is always satisfied for all $j \notin V$. Thus, to find $\alpha$ we only need to satisfy Eq. S9 for all $j$ in the violated set $V$. Now we have two sub-cases:

- If there exists $j \in V$ such that $\omega_j^{(i)} = \epsilon$, then Eq. S9 is satisfied only if $\alpha(\omega_j^{*(i)} - \omega_j^{(i)}) \geq 0$. On the other hand, because $j \in V$ and $\omega^{(i)}$ is feasible, we have $\omega_j^{*(i)} < \epsilon \leq \omega_j^{(i)} \implies \omega_j^{*(i)} - \omega_j^{(i)} < 0$. Therefore, Eq. S9 is satisfied only if $\alpha = 0$. Thus, in this case we set $\omega^{(i+1)} = \omega^{(i)}$ and add the constraint $j$ into $\mathcal{A}^{(i)}$ to obtain $\mathcal{A}^{(i+1)}$.
- Otherwise, let $\Delta_j = \frac{\omega_j^{*(i)} - \epsilon}{\omega_j^{(i)} - \epsilon}$ for all $j \in V$. After rewriting Eq. S9 and substituting $\Delta_j$ to it, we get the following condition: $\alpha \leq \frac{1}{1-\Delta_j}$ for all $j \in V$, or equivalently, $\alpha \leq \min_{j \in V} \frac{1}{1-\Delta_j} = \frac{1}{1-\Delta_p}$ where $\Delta_p$ is the minimum of all $\Delta_j$. Thus, we set $\alpha = \frac{1}{1-\Delta_p}$, $\omega^{(i+1)} = \omega^{(i)} + \alpha d$, and add $p$ into $\mathcal{A}^{(i)}$ to obtain $\mathcal{A}^{(i+1)}$.

The active-set algorithm described in this section is summarized in Algorithm 2.

#### A.2 Convergence of the EM algorithm

Recall that in the M-step, we need to find $\tau \geq 0$ and $\omega \geq \epsilon$ that satisfy $\Psi$ and maximize the following function:

$$g(\tau, \omega; q) = \sum_{i=1}^N \sum_{j=1}^k q_{ij} \log f(\hat{b}_i|\omega_j, \tau_i), \tag{S10}$$

where $f$ denote the density function of the Gaussian model for branch estimation uncertainty, as described in the main text.

Assume that the log-likelihood function $l(\tau, \omega)$ has a maximum value. At each iteration (h) of the algorithm, let $q^{(h)}$ denote the posterior computed in the E-step, $\tau(h), \omega(h)$ and $\tau(h)+1, \omega(h+1)$ denote the solution found before and after M-step. Then we claim the following:

**Claim 1.** *If $g(\tau^{(h+1)}, \omega^{(h+1)}; q^{(h)}) \geq g(\tau^{(h)}, \omega^{(h)}; q^{(h)})$, then $l(\tau^{(h+1)}, \omega^{(h+1)}) \geq l(\tau^{(h)}, \omega^{(h)})$*

*Proof.* Let $C = \sum_{i,j} q_{ij} \log(kq_{ij}^{(h)})$. Subtracting $C$ from both sides of the inequality $g(\tau^{(h+1)}, \omega^{(h+1)}; q^{(h)}) \geq g(\tau^{(h)}, \omega^{(h)}; q^{(h)})$, we have:

$$\sum_{i=1}^N \sum_{j=1}^k q_{ij}^{(h)} \log \frac{f(\hat{b}_i|\omega_j^{(h+1)}, \tau_i^{(h+1)})}{kq_{ij}^{(h)}} \geq \sum_{i=1}^N \sum_{j=1}^k q_{ij}^{(h)} \log \frac{f(\hat{b}_i|\omega_j^{(h)}, \tau_i^{(h)})}{kq_{ij}^{(h)}} \tag{S11}$$

Recall that $q_{ij}^{(h)} = \frac{f(\hat{b}_i|\omega_j^{(h)}, \tau_i^{(h)})}{\sum_{m=1}^k f(\hat{b}_i|\omega_m^{(h)}, \tau_i^{(h)})}$ and $\sum_j q_{ij}^{(h)} = 1$. Therefore, we can rewrite the right hand side (RHS) of Eq. S11 as follows

$$\begin{aligned} \text{RHS} &= \textstyle\sum_{i=1}^N \sum_{j=1}^k q_{ij}^{(h)} \log \left[\frac{1}{k} \sum_{m=1}^k f(\hat{b}_i|\omega_m^{(h)}, \tau_i^{(h)}\right] \\ &= \textstyle\sum_{i=1}^N \log \left[\frac{1}{k} \sum_{m=1}^k f(\hat{b}_i|\omega_m^{(h)}, \tau_i^{(h)}\right] \sum_{j=1}^k q_{ij}^{(h)} \\ &= \textstyle\sum_{i=1}^N \log \left[\frac{1}{k} \sum_{m=1}^k f(\hat{b}_i|\omega_m^{(h)}, \tau_i^{(h)})\right] = l(\tau^{(h)}, \omega^{(h)}) \end{aligned} \tag{S12}$$

**Algorithm 2** The active-set method to solve the problem $\mathcal{P}$ defined in Eq. S1

**function** SOLVEOMEGA($\tau, \hat{b}, \mu, \epsilon, k$)
  $\omega_j^{(1)} \leftarrow \mu$ for all $j = 1..k$ ▷ feasible initital point
  $\mathcal{A}^{(1)} \leftarrow \emptyset$
  **for** i = 1 to MaxNumberIterations **do**
    $\omega^{*(i)}, \eta^{*(i)}, \lambda^{*(i)} \leftarrow$ SOLVELAGRANGE($\mathcal{A}^{(i)}$) ▷ See Algorithm. 1
    **if** $\omega^{*(i)}$ is feasible **then**
      **if** $\lambda_j^{*(i)} \geq 0$ for every $j \in \mathcal{A}^{(i)}$ **then**
        **return** $\omega^{*(i)}$ ▷ satisfies KKT $\implies$ feasible and optimal
      **else**
        $\lambda_h^{*(i)} \leftarrow \min_{j \in \mathcal{A}^{(i)}} \lambda_j^{*(i)}$
        remove $h$ from $\mathcal{A}^{(i)}$ to get $\mathcal{A}^{(i+1)}$
    **else**
      $V = \{j | \omega_j^{*(i)} < \epsilon\}$ ▷ the violated set of $\omega^{*(i)}$
      **if** there exists $j \in V$ s.t. $\omega_j^{(i)} = \epsilon$ **then**
        add $j$ into $\mathcal{A}^{(i)}$ to get $\mathcal{A}^{(i+1)}$
        $\omega^{(i+1)} \leftarrow \omega^{(i)}$
      **else**
        $\Delta_j \leftarrow \frac{\omega_j^{*(i)} - \epsilon}{\omega_j^{(i)} - \epsilon}$ for all $j \in V$
        $\Delta_p \leftarrow \min_{j \in V} \Delta_j$
        $\alpha \leftarrow \frac{1}{1 - \Delta_p}$
        $\omega^{(i+1)} \leftarrow \omega^{(i)} + \alpha(\omega^{*(i)} - \omega^{(i)})$ ▷ feasible and "more optimal"
        add $p$ into $\mathcal{A}^{(i)}$ to get $\mathcal{A}^{(i+1)}$
  **return** the last $\omega^{(i)}$

On the other hand, applying Jensen's inequality, we get an upper bound for the left hand side (LHS) of Eq. S11:

$$\begin{aligned} \text{LHS} &\leq \sum_{i=1}^{N} \log \left[ \sum_{j=1}^{k} q_{ij}^{(h)} \frac{f(\hat{b_i} | \omega_j^{(h+1)}, \tau_i^{(h+1)})}{k q_{ij}^{(h)}} \right] \\ &= \sum_{i=1}^{N} \log \left[ \frac{1}{k} \sum_{j=1}^{k} f(\hat{b_i} | \omega_j^{(h+1)}, \tau_i^{(h+1)}) \right] = l(\tau^{(h+1)}, \omega^{(h+1)}) \end{aligned} \tag{S13}$$

Thus, from Eq. S11, Eq. S12, and Eq. S13, we have $l(\tau^{(h+1)}, \omega^{(h+1)}) \geq l(\tau^{(h)}, \omega^{(h)})$. □

**Corollary 2.** *(Claim 1 in the main text) The EM algorithm described in the main text monotonically improves the log-likelihood function after each iteration. Furthermore, if the log-likelihood $l(\tau, \omega)$ has a maximum value, then the algorithm converges.*

*Proof.* Recall that in the M-step at iteration (h) , we use coordinate descent starting at $(\tau^{(h)}, \omega^{(h)})$ to find $(\tau^{(h+1)}, \omega^{(h+1)})$. Therefore, $g(\tau^{(h+1)}, \omega^{(h+1)}; q^{(h)}) \geq g(\tau^{(h)}, \omega^{(h)}; q^{(h)})$ by construction. By Claim 1, we conclude that the algorithm monotonically improves $l$. In addition, note that $l$ has a maximum value so it is bounded above. Thus, the sequence $sn(h) = l(\tau^{(h)}, \omega^{(h)})$ is increasing and bounded above, so it converges. □

### References

[1] Jorge Nocedal and Stephen J. Wright. *Numerical Optimization.* Springer, New York, NY, USA, 2e edition, 2006.

### B Supplementary figures and tables

Table S1: Parameters and statistics of the 12 clock models. Lognormal distributions are parameterized by $\mu$ and $\sigma$, which are the *actual* mean and standard deviation of the distribution. The bimodal, trimodal, and quartmodal distributions are mixtures of $2, 3$, or 4 Lognormal distributions, respectively, and $p_i$ is the probability mass of component $i$ of the mixture. Gamma distribution is parameterized by its shape $\alpha$ and rate $\beta$. The other distributions are shown by their canonical parameterization.

| Model name | Parameters | Mean | Std | CV | Newly simulated |
|---|---|---|---|---|---|
| LogNormal | $\mu = 0.006, \sigma = 0.0024$ | 0.006 | 0.0024 | 0.4 | No |
| Gamma | $\alpha = 6.05, \beta = \alpha/\mu$ where $\mu = 0.006$ | 0.006 | 0.00244 | 0.407 | No |
| Exponential | $\lambda = 1/\mu$ where $\mu = 0.006$ | 0.006 | 0.006 | 1.0 | No |
| Uniform | $a = 0, b = 0.012$ | 0.006 | 0.0035 | 0.577 | Yes |
| Bimodal 1 | $\mu_1 = 0.003, \sigma_1 = 0.0003, p_1 = 0.5$<br>$\mu_2 = 0.009, \sigma_2 = 0.0003, p_2 = 0.5$ | 0.006 | 0.003 | 0.5 | Yes |
| Bimodal 2 | $\mu_1 = 0.002, \sigma_1 = 0.0003, p_1 = 0.5$<br>$\mu_2 = 0.01, \sigma_2 = 0.0003, p_2 = 0.5$ | 0.006 | 0.004 | 0.667 | Yes |
| Bimodal 3 | $\mu_1 = 0.003, \sigma_1 = 0.0024, p_1 = 0.5$<br>$\mu_2 = 0.009, \sigma_2 = 0.0024, p_2 = 0.5$ | 0.006 | 0.0038 | 0.641 | Yes |
| Bimodal 4 | $\mu_1 = 0.002, \sigma_1 = 0.0024, p_1 = 0.5$<br>$\mu_2 = 0.01, \sigma_2 = 0.0024, p_2 = 0.5$ | 0.006 | 0.0047 | 0.783 | Yes |
| Trimodal 1 | $\mu_1 = 0.002, \sigma_1 = 0.0003, p_1 = 0.2$<br>$\mu_2 = 0.006, \sigma_2 = 0.0003, p_2 = 0.6$<br>$\mu_3 = 0.01, \sigma_3 = 0.0003, p_3 = 0.2$ | 0.006 | 0.00254 | 0.423 | Yes |
| Trimodal 2 | $\mu_1 = 0.002, \sigma_1 = 0.0003, p_1 = 0.4$<br>$\mu_2 = 0.006, \sigma_2 = 0.0003, p_2 = 0.2$<br>$\mu_3 = 0.01, \sigma_3 = 0.0003, p_3 = 0.4$ | 0.006 | 0.0036 | 0.6 | Yes |
| Trimodal 3 | $\mu_1 = 0.002, \sigma_1 = 0.0003, p_1 = 0.333$<br>$\mu_2 = 0.006, \sigma_2 = 0.0003, p_2 = 0.333$<br>$\mu_3 = 0.01, \sigma_3 = 0.0003, p_3 = 0.333$ | 0.006 | 0.003 | 0.5 | Yes |
| Quartmodal | $\mu_1 = 0.001, \sigma_1 = 0.0003, p_1 = 0.25$<br>$\mu_2 = 0.004, \sigma_2 = 0.0003, p_2 = 0.25$<br>$\mu_3 = 0.008, \sigma_3 = 0.0003, p_3 = 0.25$<br>$\mu_4 = 0.011, \sigma_4 = 0.0003, p_4 = 0.25$ | 0.006 | 0.0038 | 0.633 | Yes |

Table S2: The 5 scenarios of the simulated angiosperms

| Scenario | Description |
|---|---|
| Scenario 1 | herbaceous : woody angiosperm = 3:1 |
| Scenario 2 | herbaceous : woody angiosperm = 6:1 |
| Scenario 3 | angiosperm : gymnosperm = 4:1<br>herbaceous : woody angiosperm = 3:1 |
| Scenario 4 | angiosperm : gymnosperm = 4:1<br>herbaceous : woody angiosperm = 3:1<br>Gnetales take the rates of herbaceous angiosperm |
| Scenario 5 | angiosperm : gymnosperm = 4:1<br>herbaceous : woody angiosperm = 3:1<br>Gnetales take the rates of woody angiosperm |

Table S3: The estimated divergence time of the HIV-1 subtypes

| Subtype | MD-Cat | Wertheim 2012 | Patino 2017 | Abecasis 2009 |
|---|---|---|---|---|
| A1 | 1950 | 1952 (1947-58) | 1953 (1944–60) | 1955 (1940-1969) |
| B | 1950 | 1959 (1953-63) | —— | 1959 (1951-1967) |
| C | 1946 | 1947 (1940-54) | —— | 1952 (1940-1962) |
| D | 1945 | 1952 (1946-56) | 1956 (1927–61) | 1944 (1935-1952) |
| F1 | 1960 | 1955 (1948-61) | —— | —— |
| F2 | 1963 | 1966 (1961-70) | —— | —— |
| G | 1954 | —— | 1969 (1961–74) | 1969 (1960-1977) |

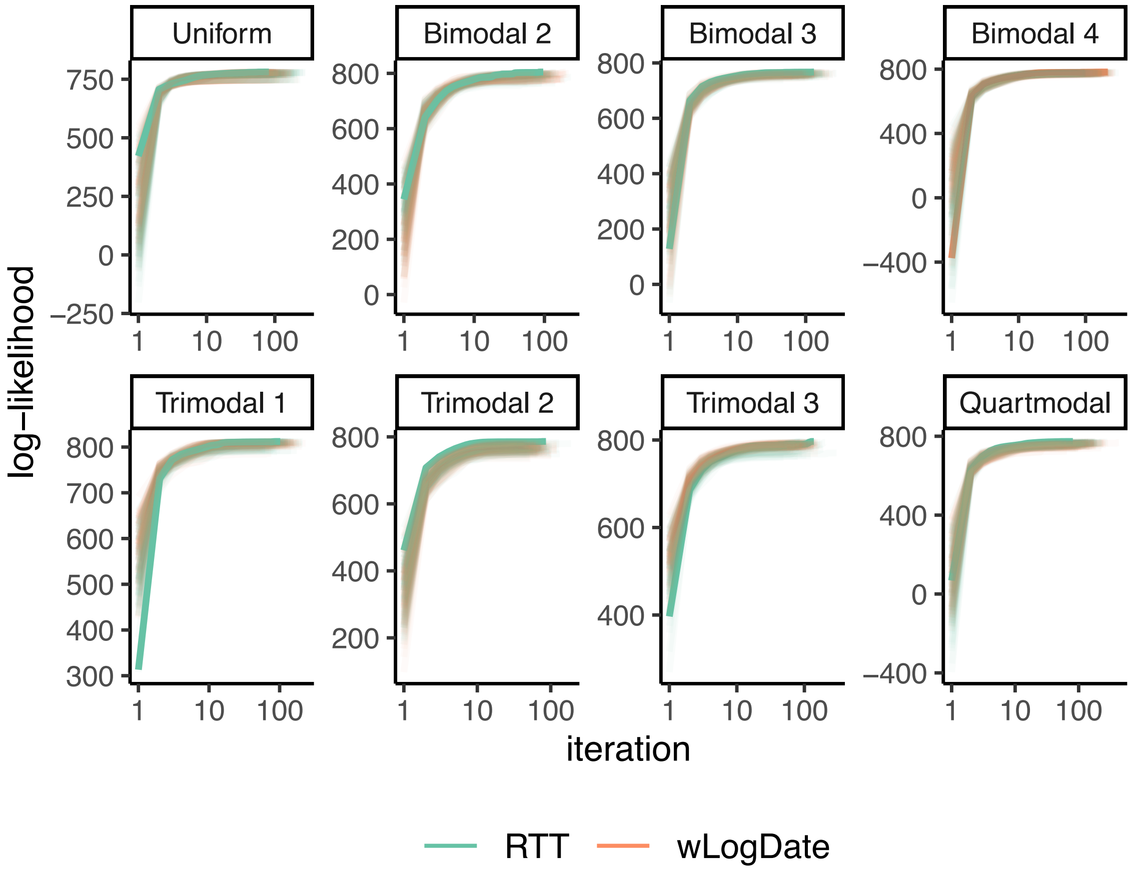

Figure S1: Log-likelihood versus iterations of the EM algorithm on the first replicate using 200 initial points. The thick line shows the best final log-likelihood.

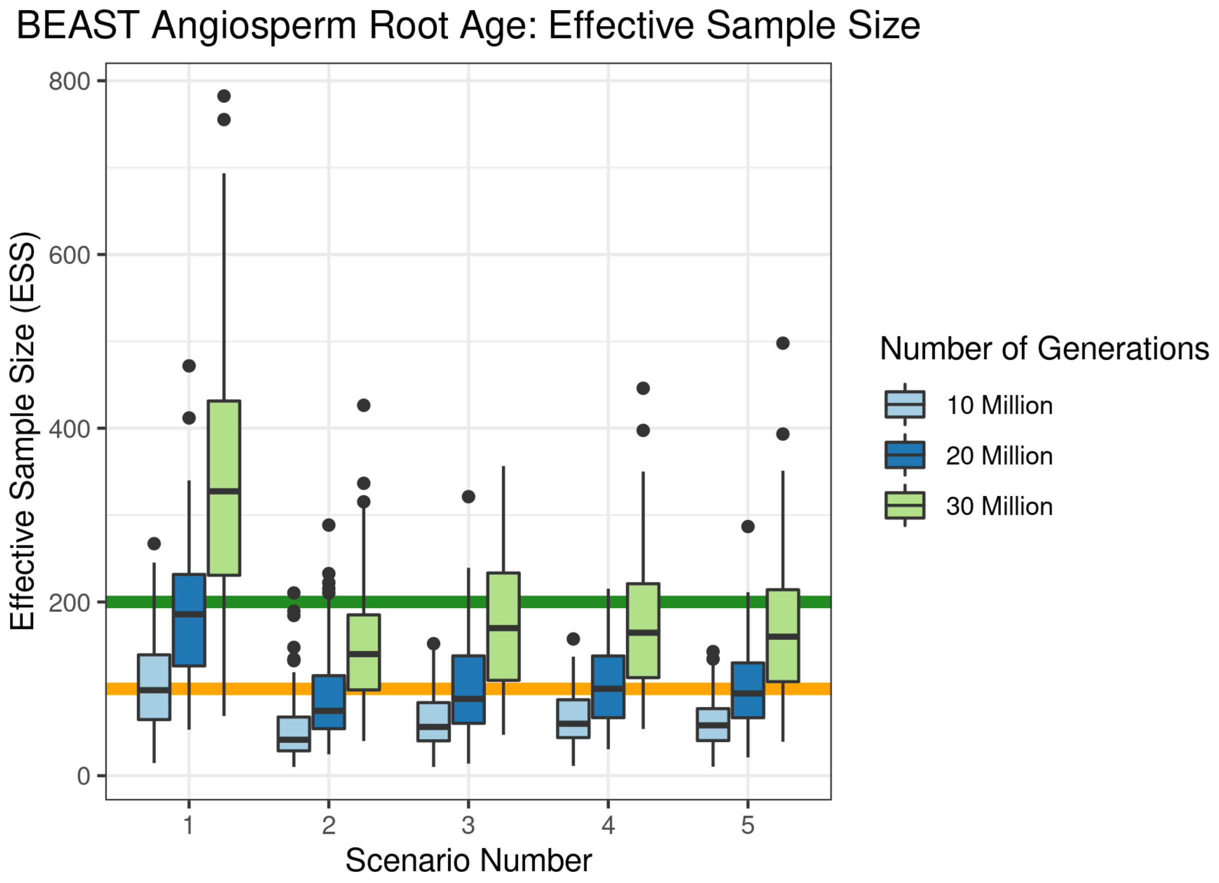

Figure S2: Effective sample size (ESS) of the posterior for root age in the 5 different angiosperm tree scenarios when analyzed on BEAST. Different values are shown when analysis was run for 10, 20, and 30 million generations. ESS above the yellow horizontal line (y = 100) report the minimum required to demonstrate MCMC convergence, while those above the green line (y = 200) are more clearly converged.

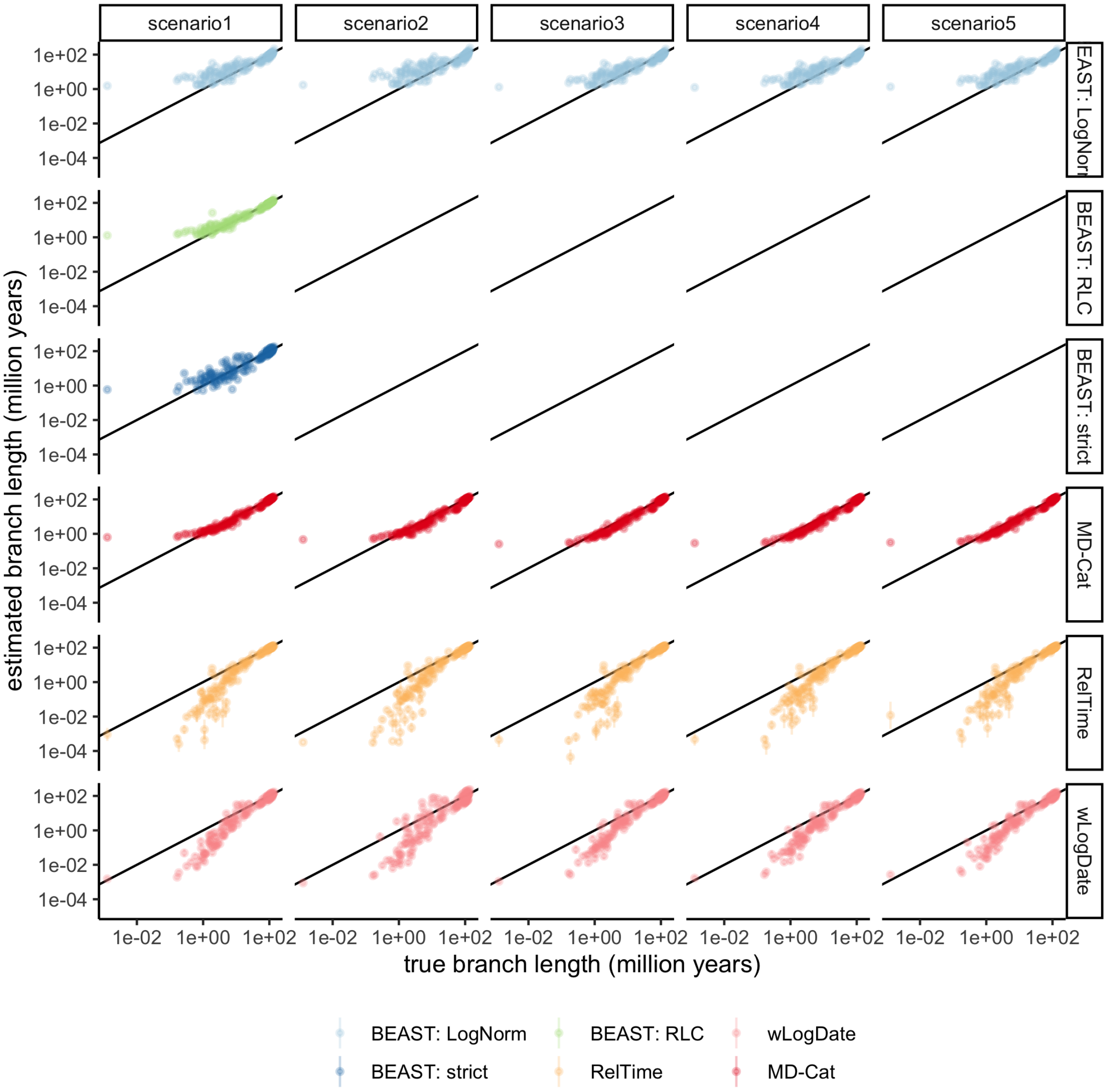

Figure S3: Estimated versus true branch length. Each dot shows the mean and standard error over all 100 replicates for that branch in the true tree. Both axis are in log scale.

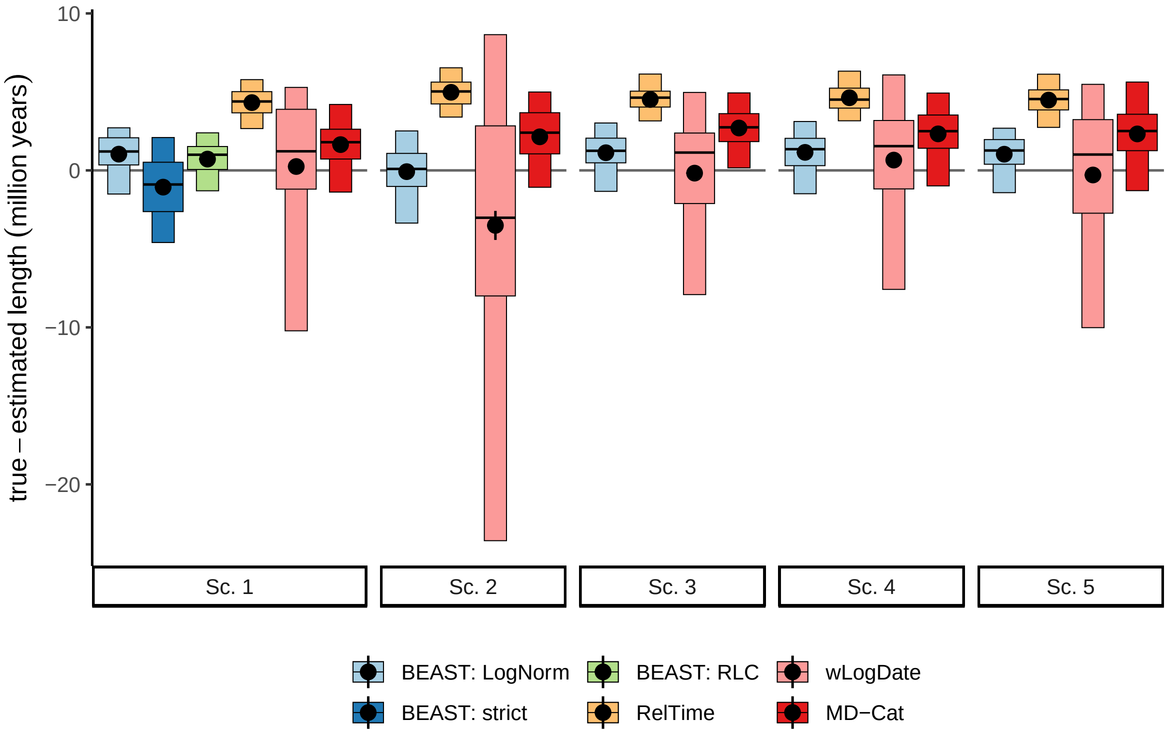

Figure S4: The distribution of the mean difference between true and estimated branch length across all branches of each replicate, indicating bias. We show mean (black dots), and standard error, 25-75% percentiles (wide boxes), and 5-95% percentiles (narrow boxes) over 100 replicates.

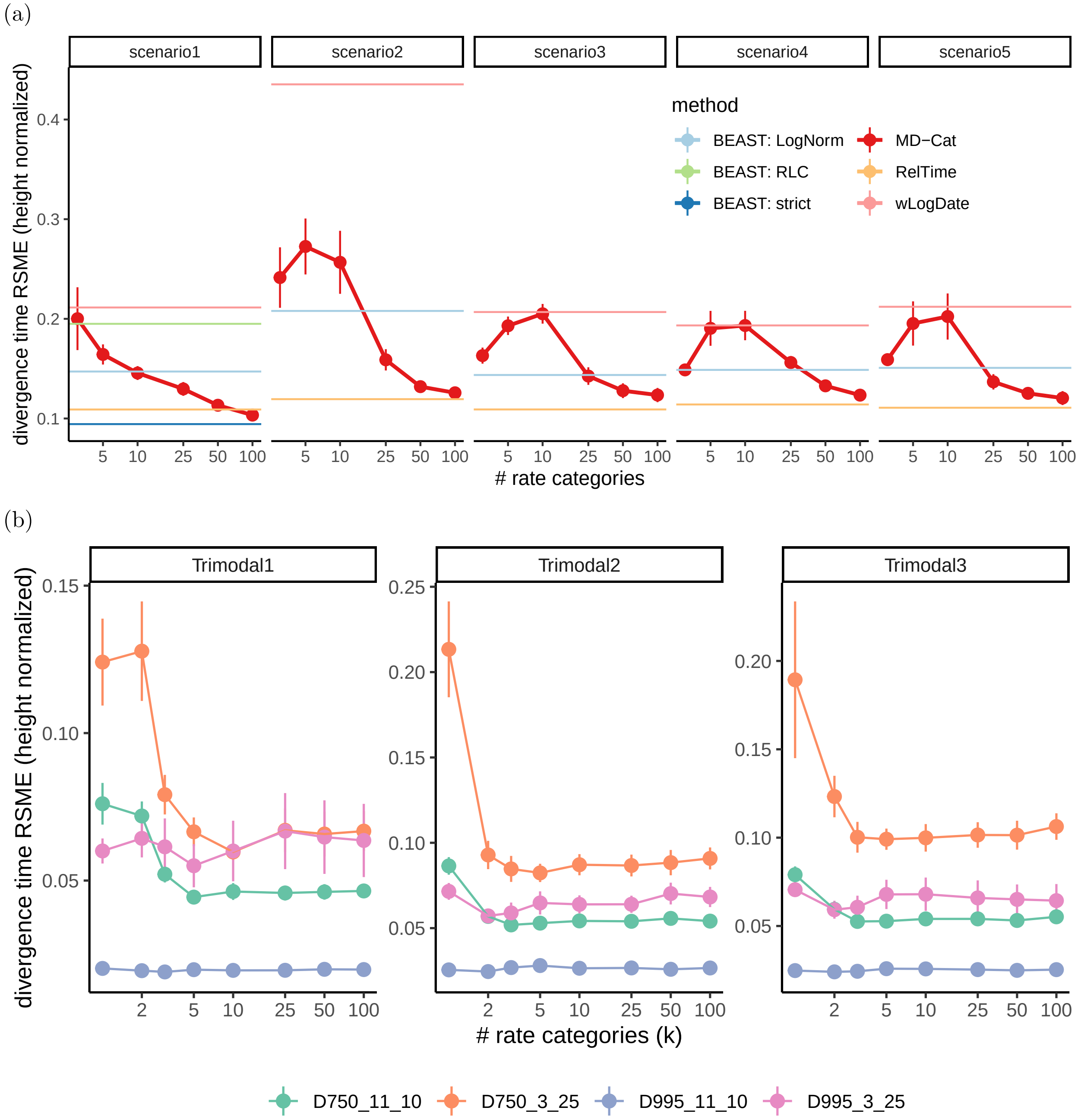

Figure S5: Analysis of the sensitivity of MD-Cat to the number of rate categories ($k$) for (a) the Angiosperms simulated data, and (b) the HIV simulated dataset under the three trimodal clock models. The default $k$ used in all other analyses is 50. We compare methods in terms of the divergence time error, defined as the root mean squared error of divergence times of all nodes versus true divergence, normalized by tree height.

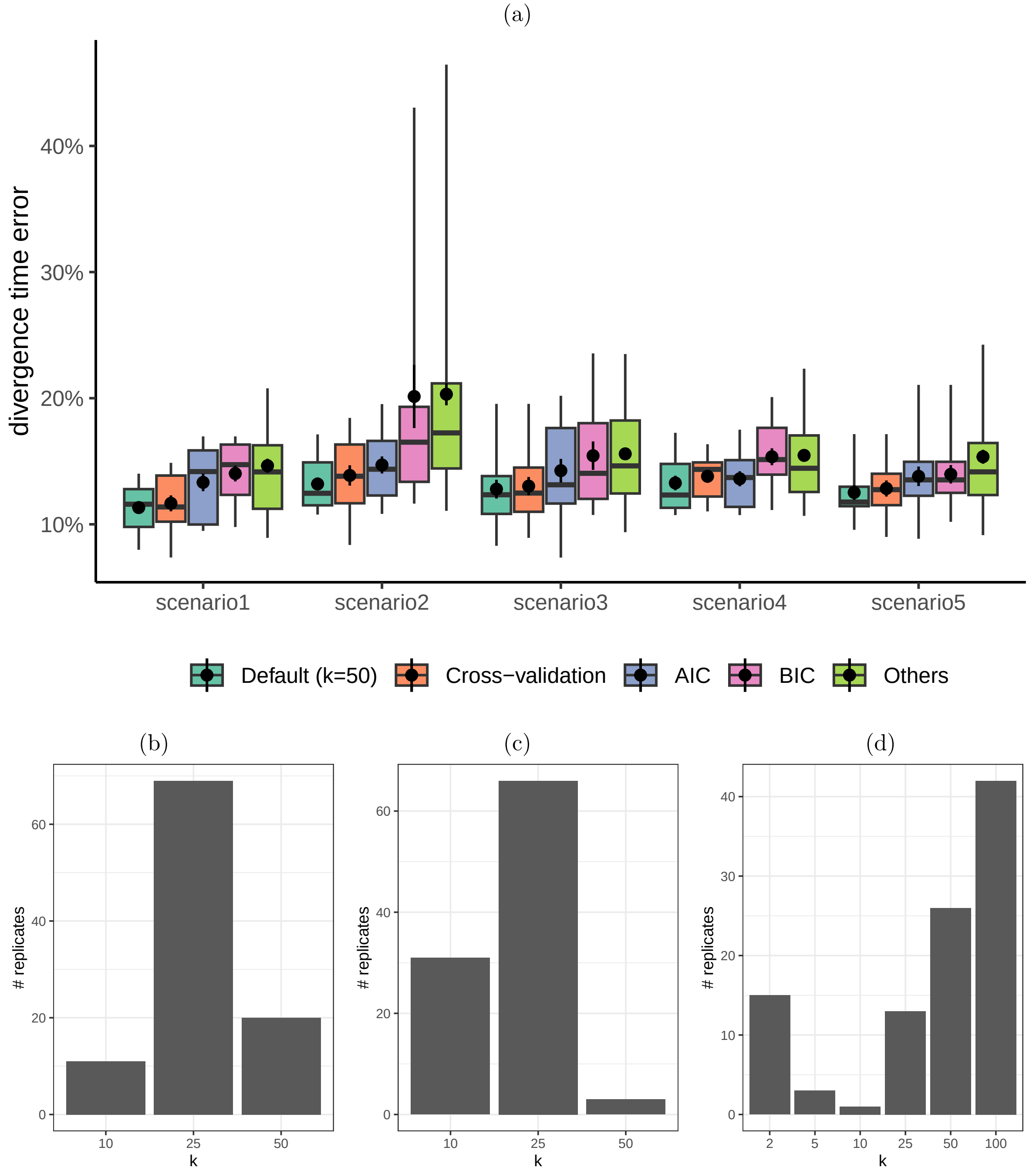

Figure S6: Selection on $k$ for the Angiosperms simulated data using AIC, BIC, and cross-validation. (a): The divergence time error (absolute difference of true and estimated normalized by tree height) across all branches. b,c,d: The selected $k$ distribution across all runs for AIC, BIC, and cross-validation, respectively.

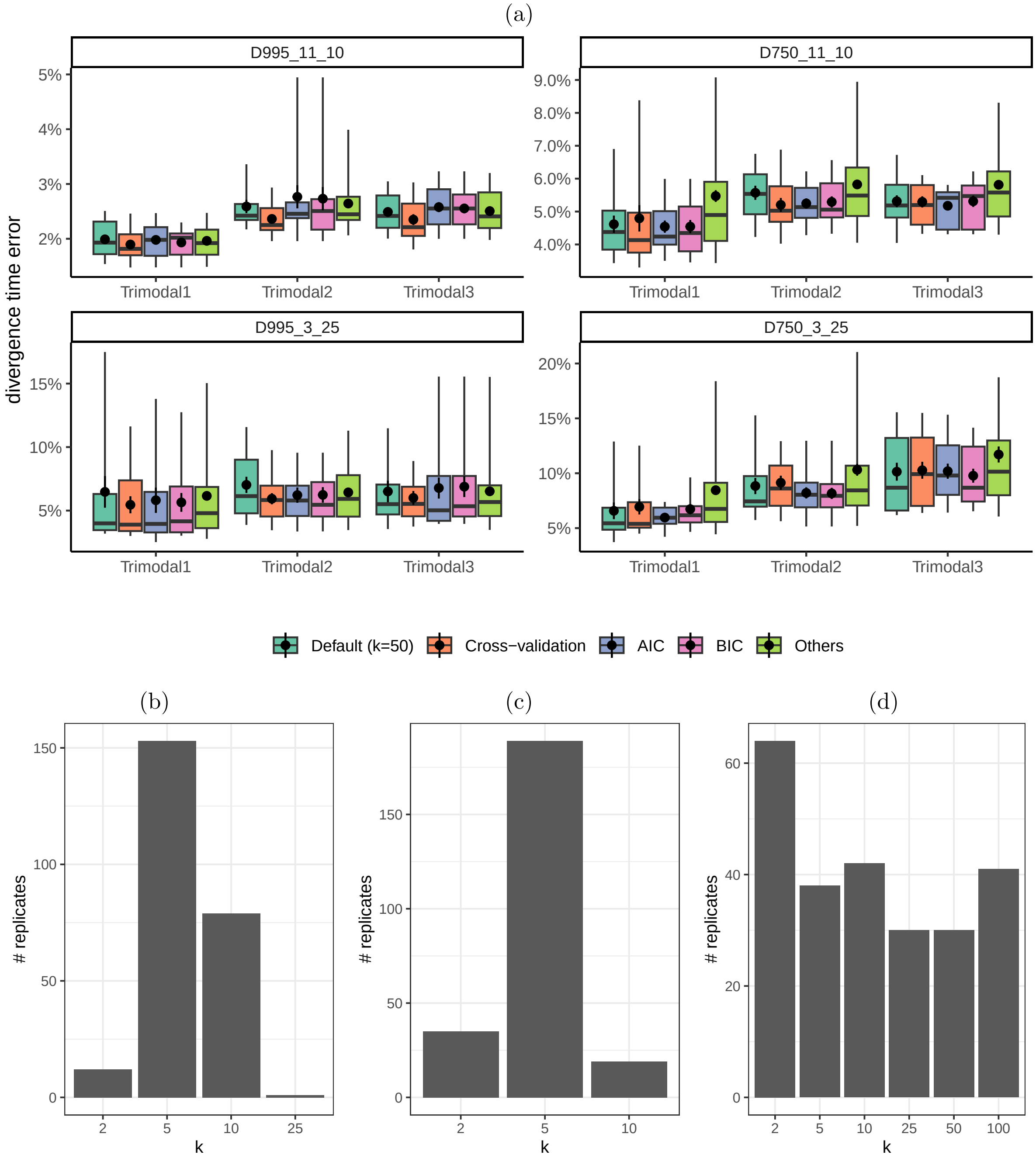

Figure S7: Selection on $k$ for the HIV simulated data using AIC, BIC, and cross-validation. (a): The divergence time error (absolute difference of true and estimated normalized by tree height) across all branches. b,c,d: The selected $k$ distribution across all runs for AIC, BIC, and cross-validation, respectively.

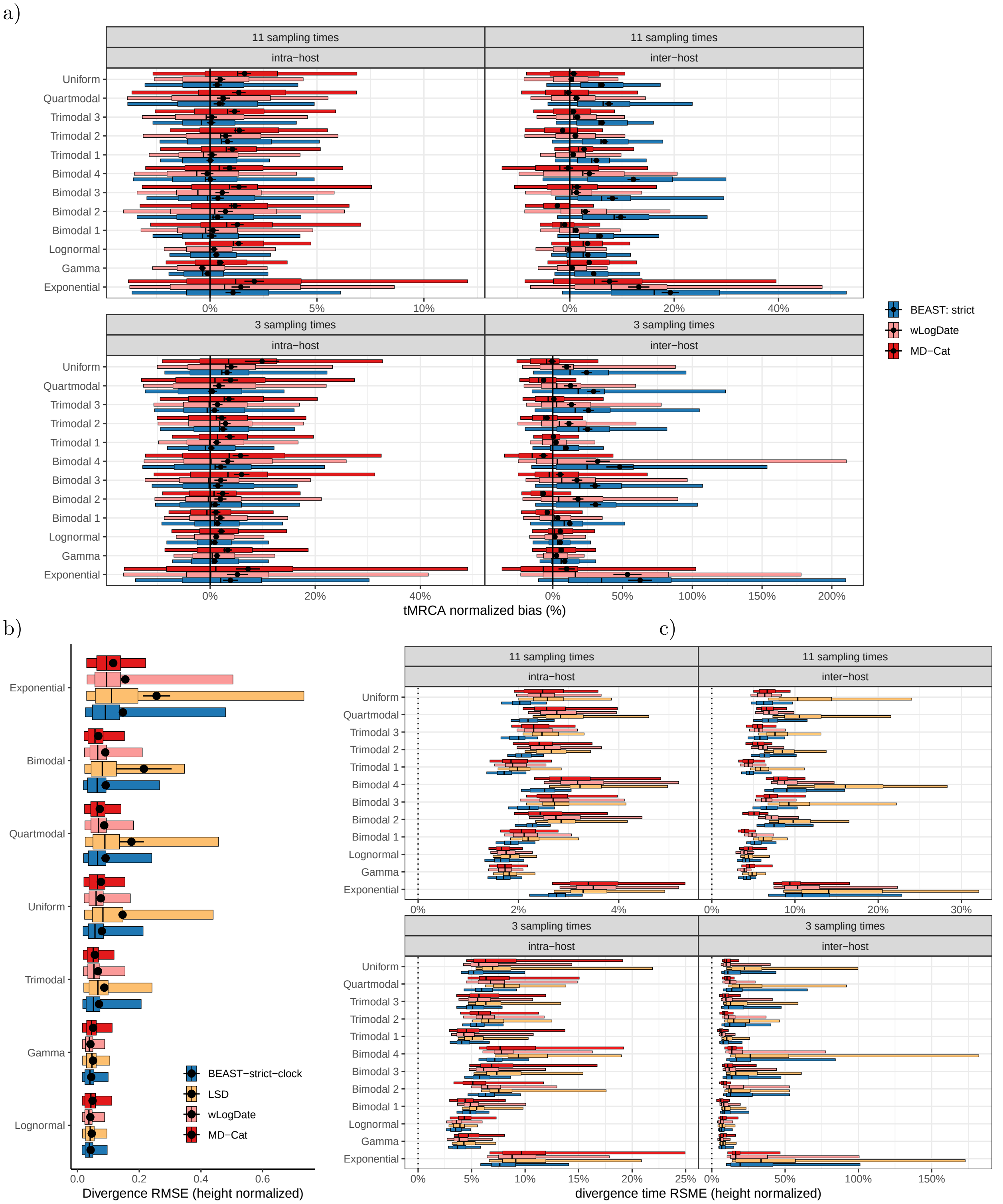

Figure S8: Detailed comparisons on the HIV simulated dataset. a) Normalized tMRCA error, broken down by tree condition and clock model. Since LSD can have extremely high errors, we removed it from the figure to help readability. Narrow boxes show 5-95 percentiles and wide boxes show 25-75 percentiles. Bar shows median. Mean and standard error are also shown as dots and error bars. b) Divergence time RMSE (normalized by tree height). Mean/se omitted due to high mean error for LSD. Gamma and Lognormal are unimodal. Uniform and Exponential are nonmodal. c) Same as (b) with tree models and clock models separated.

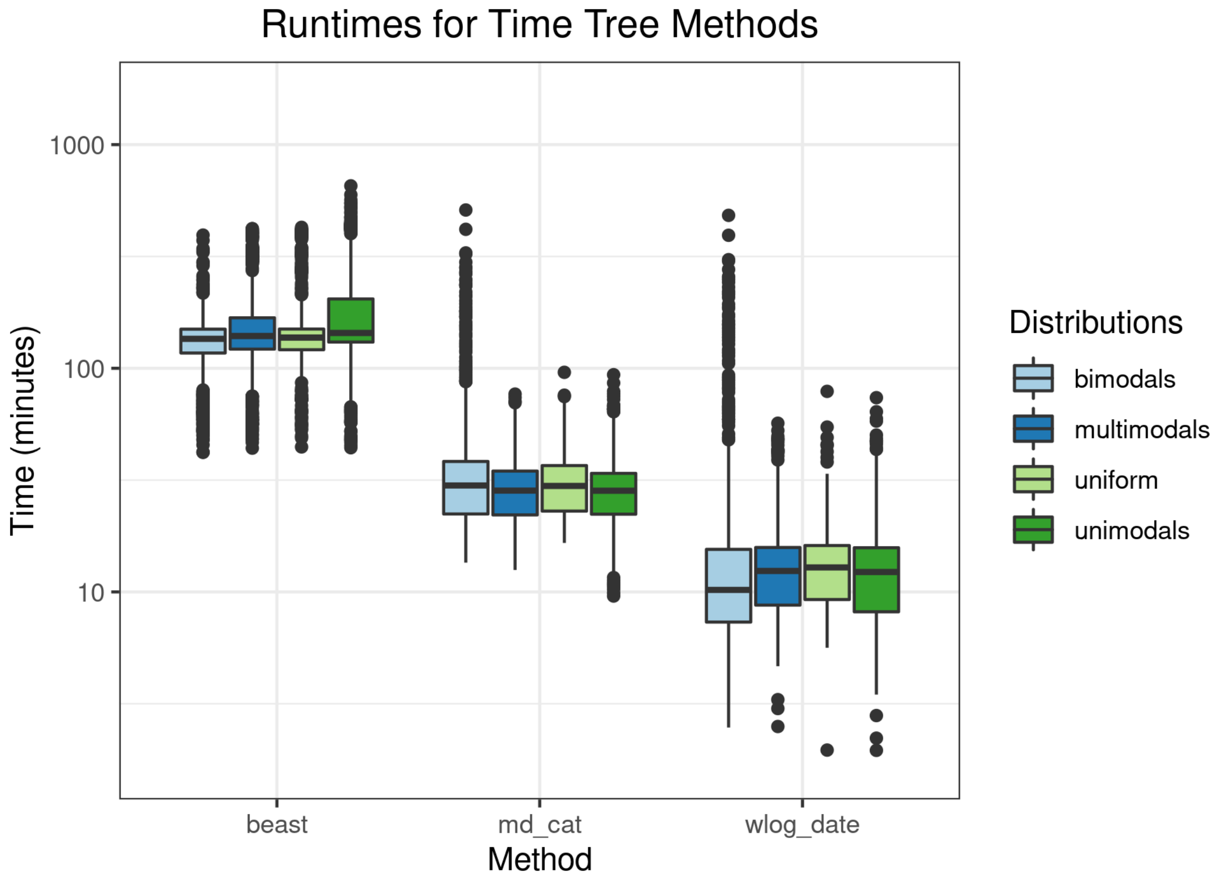

Figure S9: Comparison of the running time of BEAST, MD-Cat, and wLogDate separated into four different clock model groups. The y-axis represents the time in minutes and is shown in the log-scale. All analyses were run on Intel Xeon CPU E5-2670 (103,597,240 kB of memory). MD-Cat runtime does not include CI calculation and SU branch length estimation but those would add a negligible amount: about 2 minutes for CI estimation and several seconds for SU branch length estimation using RAxML.

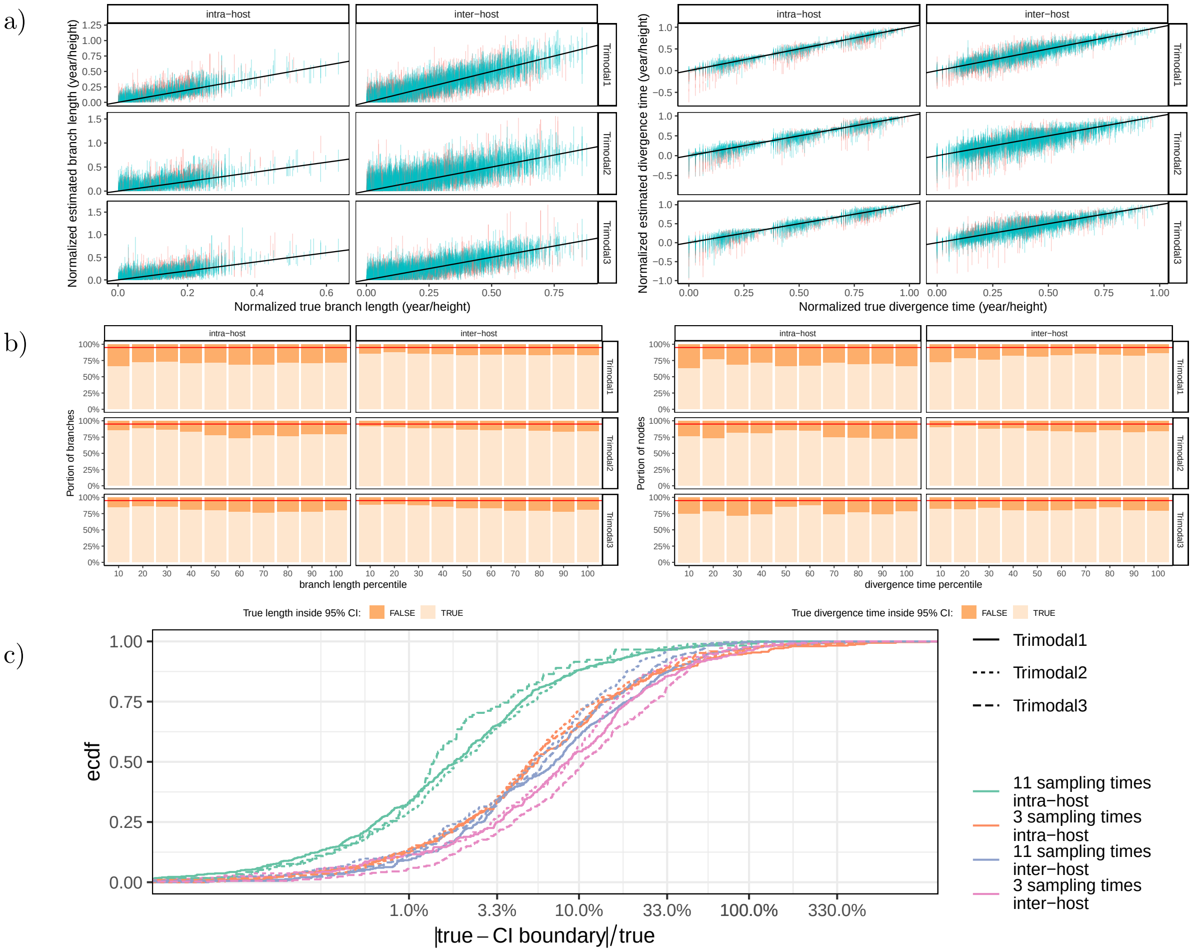

Figure S10: Evaluating support values. a) Confidence interval (CI) estimated by MD-Cat for branch lengths (Left) and divergence times (Right). Blue: CI includes the true value; red: it does not. b) The portion of branches (Left) or nodes (Right) that fall outside the 95% CI, shown separately for 10 deciles of true branch length / divergence time. c) For nodes where the true value falls out of CI, we show how far it falls as a percentage. We show the distribution of $|t - c|/t$ where $c$ is the CI boundary which fails to capture the true value and $t$ is the true divergence time.
